# On the applicability domain of HADDOCK3 for protein-aptamer docking: documented failure modes from a 5×7 cross-target screening matrix and a 1676 aa receptor case study (P01031)

**DOI:** 10.64898/2026.05.11.724398

**Authors:** Eisuke Dohi

## Abstract

We screened a 5-receptor x 7-aptamer = 35-cell cross-target screening matrix with HADDOCK3 under blind ambiguous-interaction-restraint (AIR) protocols on AlphaFold-modelled receptors. **The 35-cell matrix is primarily a cross-target / decoy screening matrix rather than a 35-cognate-pair benchmark:** it contains an n = 4 K_D-calibration subset under matched assay conditions, at least six biological cognate or intended-cognate cells, and the remaining cells are intentional non-target pairings used to characterise score-distribution behaviour. The screen surfaced 12 operationally distinct failure modes that collapse into five broad conceptual groups. The principal case study is **P01031** (complement C5, 1676 aa, ≥ 12 structural domains): all seven panel members produced positive HADDOCK3 top-1 scores under a scale-adaptive AIR. Score-term decomposition locates the anomaly in the AIR term (+217 to +268 to top-1 score). With AIR zeroed, scores fall to -131 to -74 -- the small-receptor regime. Boltz-2 cofolding chain-pair ipTM (cpi_AB) is an independent channel: P01031 shows the lowest median cpi_AB (0.211; 0/7 above the 0.5 confident-interface threshold). To our knowledge, this is an early documented case study of a 1676 aa multi-domain receptor exhibiting this signature under a blind scale-adaptive AIR workflow -- an n = 1 mechanistic case, not a statistical generalisation. We adapt the QSAR *applicability-domain* concept to in silico aptamer screening. We report an empirical Mode 1 mitigation, a pLDDT-aware AIR prefilter, with cohort Jaccard recovery of ∼10x. The n = 4 K_D-calibration Spearman ρ shift is reported as exploratory cross-method convergence, not as a calibration claim.

## 1. Introduction

HADDOCK3 [1] is the most recent major release of the HADDOCK family of physics-based, restraint-driven dockers, succeeding the widely-deployed HADDOCK2.4 web server [2] which has served over half a million docking jobs since 2008. Three extensions have appeared in 2024-2026. First, MARTINI2 coarse-graining now scales to a ∼6 kAa PFK filament (PDB 8W2G/8W2I/8W2L; ≈ 6240 aa across 8 chains; exact residue count not stated in [3]). Second, AlphaFlow + HADDOCK addresses antibody H3-loop ensembles [4]. Third, protein-DNA / protein-NA benchmarking has been published on systems such as ComP + dsDNA Uptake Sequence [5]. The CAPRI rounds 47-55 established the AI-derived contact-prior paradigm for AIRs [6], with the operational recipe in [7]: AlphaFold-derived contact predictions are converted to HADDOCK ambiguous restraints using residue-level confidence thresholding. Recent generative-prior work [8, 9] extends the AI-prior toolkit further upstream.

For *aptamer*-protein docking specifically, the published benchmark coverage is narrower than is sometimes assumed. In the literature we reviewed, we did not identify a published HADDOCK3, AlphaFold-3 [10], or cofolder benchmark -- including RoseTTAFold2NA [11] and Chai-1 [12] -- that covers the ≥ 1500 aa receptor + nucleic-acid ligand regime. Independently, the AF3-RNA benchmark of Bernard et al. [13] documents that AF3 lags human-aided methods on CASP-RNA and fails to generalise to orphan RNA families without context (INF-nWC < 0.5 across five RNA test sets), motivating caution at the protein-aptamer interface as well. The HADDOCK3 reference paper [1] validates antibody-antigen, glycan, scoring, and alascan use cases (no protein-NA test among the four). The CAPRI-rounds paper [6] states no specific receptor-size envelope. The ComP study [5] used a 136 aa receptor. The third-party AF3 protein-NA benchmark of Peng et al. (Benchmark-IX) [14] was restricted to *“complexes with total residue count < 1000”* (page 3 verbatim).

The most directly comparable cofolder-only benchmark is Zhao et al. [15]: it evaluates AlphaFold3, Chai-1, Boltz-2, and RoseTTAFold2NA on 11 protein-aptamer complexes (curated from PDB entries released after the AF3 training cutoff of 30 September 2021; Methods §Datasets and Table 2 of [15]), with explicit caution that Boltz-2 ipTM can be overconfident on protein-aptamer systems. Crucially, Zhao et al. do not evaluate HADDOCK3 and therefore do not address the AIR-term artefacts catalogued here.

Cataloguing the failure modes of a docking protocol is methodologically valuable for three reasons. First, it defines an applicability domain -- the set of (target, ligand, model-state) tuples on which the protocol is validated to work -- by analogy with the QSAR applicability-domain concept [16, 17, 18]. Second, it separates protocol-fixable issues (e.g., AIR specification) from method-fundamental ones (e.g., affinity-head training-distribution restrictions). Third, it enables transparent reporting of where a pipeline should and should not be deployed before any wet experimental commitment.

We present a catalogue of 12 failure modes encountered during a 5-receptor x 7-aptamer = 35-cell cross-target screening matrix. The K_D-calibration subset is n = 4 cells with literature K_D records under matched assay conditions (TBA-thrombin, Pegaptanib-VEGFA, AX102-PDGFB, S10yh2-ITGB3). The broader cohort includes ≥ 6 biological cognate or intended-cognate cells. HD22-thrombin, RE31-thrombin, and Avacincaptad-C5 are intended cognates without matched-condition K_D records. S10yh2 is a partial cognate because P05106 is modelled as the β3 monomer rather than the αIIbβ3 heterodimer (Methods §2). The catalogue is complementary to Zhao et al. [15]: where they report a cofolder-only benchmark, we report HADDOCK3-specific failure modes plus an empirical AIR-feed mitigation. A separate companion manuscript will report the broader multi-method validation cohort. The present manuscript is self-contained with respect to the 35-cell failure-mode catalogue, the P01031 case study, and the pLDDT-aware AIR mitigation analysis.

This paper makes four contributions:

1. A 12-mode failure catalogue with severity scoring, raw evidence per mode, and conceptual collapse to five broad mode classes (Classes A-E; Table 1, §3.1, Figure 1).
2. An empirical case study on P01031 (1676 aa) showing 7/7 positive, AIR-penalty-dominated HADDOCK3 scores compared against Boltz-2 cofolding (Figure 2).
3. An applicability-domain map placing P01031 outside the approximate published HADDOCK3 / cofolder NA-ligand benchmark coverage region (Figure 3).
4. An empirical Mode 1 mitigation (Figure 4; §3.7), including cohort-level Jaccard recovery of ∼10x; a per-receptor floor/cap engagement analysis, a random-baseline correction showing that the baseline failure is ∼40x below random expectation, an exploratory n = 4 K_D-calibration Spearman ρ analysis (exact two-sided p = 0.33; treated as cross-method convergence, not calibration), and a DockQ benchmark on four reference complexes with an explicit PDB-memorisation caveat.

**Table 1.** Twelve documented operational failure modes of HADDOCK3 + AlphaFold-derived blind AIR for aptamer-protein docking. Severity = high / medium / low. Dataset codes: COH = present cohort (this work); LIT = literature anchor; PAR = parallel cohort analysis. Evidence basis indicates whether the mode is observed (this cohort), parallel cohort analysis, literature-supported, or extrapolated. Mitigation tested: = empirically verified in this cohort; = published in cited reference; -- = not yet tested.

| # | Mode | Axis | Class | Receptor / aptamer where observed | Severity | Evidence basis | Dataset | Mitigation | Tested? | Effective? |
| --- | --- | --- | --- | --- | --- | --- | --- | --- | --- | --- |
| 1 | Disordered tail dominates AIR | AIR | A | P00734 thrombin (residues 1-315 captured, catalytic domain missed); generalises across cohort | high | observed (this cohort) | COH, PAR | protocol-fix (pLDDT $\geq 70$ prefilter) | ✓ | yes (Jaccard $\sim 10\times$, §3.7) |
| 2 | Multi-domain receptor scatter ($\geq 1500$ aa) | AIR / receptor | B | P01031 C5 (12+ domains, 80 active over non-overlapping surfaces) | high | observed (this cohort) | COH | applicability-domain | -- | n/a (case study) |
| 3 | Fixed N = 30 distorts mass across cohort | AIR | A | all 5 receptors at fixed N = 30 (variant A baseline) | medium | observed (this cohort) | COH, PAR | protocol-fix (scale-adaptive N_active) | ✓ | yes (§3.7.2) |
| 4 | Sparse blind AIR fails (opposite end) | AIR / method | D | Giulini 2025 IL-1β / gevokizumab Use Case 1 (PDB 4G6M): sparse-AIR run yields top-ranked DockQ 0.034, consistent with under-restraint [1] | medium | literature-supported | LIT | applicability-domain | ✓ | n/a (literature) |
| 5 | Cβ-neighbor heuristic ≠ binding pocket | AIR | A | all 5 receptors (anti-correlated with literature epitopes) | high | observed (this cohort) | COH, PAR | protocol-fix (pLDDT prefilter + literature anchor when available) | ✓ | yes (parallel cohort analysis) |
| 6 | G-quadruplex aptamer geometry not predictable de novo | aptamer | C | TBA (15-nt) G4 self-fold not recovered by cofolders without templating; Stafflinger 2025 NAR Breakthrough article 9HRF/9HRD/9HRG [33] | high | literature-supported | LIT | method-fundamental (templated structure or experimental input) | ✓ | n/a |
| 7 | TBA G4 self-fold dominates cofolder ipTM | aptamer | C | TBA x multiple targets in SELEX-cross panel | medium | parallel cohort analysis | PAR | applicability-domain (per-target z-score normalisation) | ? | partial; reported separately |
| 8 | Apo-state induced-fit drift (cofolders) | aptamer / receptor | D | extrapolated from kinase systems [34] (~ 40 % Apo Drift) | medium | extrapolated | LIT | method-fundamental (multi-conformation ensemble) | -- | n/a |
| 9 | Receptor ≥ 1500 aa unvalidated for NA | receptor | B | P01031 C5 (1676 aa); Peng 2025 cap < 1000 residues total [14] | high | observed (this cohort) | COH, LIT | applicability-domain | -- | n/a (case study) |
| 10 | Multi-conformation receptors with large interdomain rearrangements (C3/C5 family) | receptor | D | C3 / C5 family [35, 36] | medium | literature-supported | LIT | method-fundamental (ensemble docking) | -- | n/a |
| 11 | Boltz-2 affinity head not validated for NA | method-maturity | D | 34/34 attempts failed schema validation | high | observed (this cohort) | COH | method-fundamental | ? | n/a (confirmed unavailable) |
| 12 | No co-structure anchor for FDA-approved Avacincaptad-C5 | method-maturity | E | Avacincaptad pegol x C5 (no PDB deposit as of 2026-05; ARVO 2025 abstract only [37, 38]) | medium | literature-supported | LIT | applicability-domain (negative observation) | -- | n/a |

### 1.1 Closely related work and differentiation

This section places three recent lines of work in direct relation to the present manuscript and states our incremental contribution against them.

Williams et al. [19] categorise low-pLDDT AlphaFold2 regions into three classes: *near-predictive* (high packing, low outliers), *pseudostructure* (low packing, low outliers, misleading SS-like appearance), and *barbed wire* (low packing, high outliers, wide looping coils). The disordered-terminus pattern that dominates the variant-A blind-AIR active-residue selections in our cohort (97 % cohort median, §3.7.4) corresponds operationally to the *barbed wire* class. We extend the Williams taxonomy into a downstream restraint-driven docking context.

Yu et al. [20] report that AlphaFold2 predicts residue-level solvent accessibility with high accuracy across taxa (Pearson r ≈ 0.815 against reference accessibility on eukaryotic, bacterial, and archaeal proteomes). They do not examine the failure of Cβ-neighbor blind heuristics on disordered termini. Their result nonetheless motivates the *mitigation* in §3.7: because high-pLDDT regions carry reliable surface-accessibility information, a pLDDT ≥ 70 prefilter recovers the binding-competent surface envelope.

Holcomb et al. [21] document an analogous AlphaFold-receptor docking bias for protein-ligand AutoDock-GPU: 41 % vs 17 % success at ≤ 2 Å RMSD on PDBbind n = 2,474 when the same ligand library is docked against crystal vs AlphaFold-2 receptors. §3.7 documents an early protein-nucleic-acid HADDOCK3 analogue of that bias category at the AIR-feed step; the Holcomb statistic itself is referenced (not re-stated) in §3.7.

Our contribution is therefore *not* the underlying mechanism. Low-pLDDT regions are well known to be unreliable, and the AlphaFold-DB FAQ recommends pLDDT ≥ 70 for downstream use. Our contribution is instead the systematic quantification of the failure mode for a specific tool combination -- *AlphaFold receptor x HADDOCK3 x Cβ-blind active-residue heuristic x protein-aptamer ligand* -- across a 5 x 7 = 35-cell cross-target matrix. We add a per-receptor floor/cap engagement analysis showing that the operational difference between fixed-N and scale-adaptive prefilters exceeds the cohort-mean figure. We also add a random-baseline correction quantifying the failure at ∼40x below random expectation. Recent cofolder benchmarks [14, 15, 22] and training-memorisation analyses [23] ask which cofolder predicts the bound complex best; we instead ask how the AIR feed should be constructed when an AlphaFold receptor is fed into HADDOCK3 -- a complementary question. Xu, Giulini, Bonvin [4] address low-pLDDT artefacts by clustering AlphaFlow generative ensembles before HADDOCK, a per-target, computationally intensive procedure. The single-pass pLDDT-filtered AIR strategy in §3.7 is a lightweight alternative for high-throughput screening.

## 2. Methods

### 2.1 Cross-target screening matrix

Five receptors spanning a ∼7x size range and contrasting topologies were selected:

- **P00734** -- thrombin, 622 aa.
- **P01127** -- PDGF-B (PDGFB chain of PDGF-BB), 241 aa.
- **P15692** -- VEGFA (Vascular Endothelial Growth Factor A), **395 aa** (AlphaFold-DB v4 model AF-P15692-F1-model_v4 used as input to HADDOCK3; this model is isoform-derived and differs from the canonical UniProt P15692-1 pre-pro form, 412 aa, and the secreted VEGF165 mature form, 165 aa).
- **P05106** -- ITGB3 (integrin β3 subunit, 788 aa, single chain). The functional binding partner of integrin-targeting aptamers is the αIIbβ3 heterodimer formed with ITGA2B (αIIb, P08514, 1039 aa); the heterodimer is not modelled in this study (limitation noted; full αIIbβ3 cryo-EM is PDB 8T2U / 8T2V) [24].
- **P01031** -- complement C5, 1676 aa, multi-domain.

Receptor models were taken from AlphaFold-DB v4 EMBL-EBI deposits (AlphaFold 2 architecture [25]; full-length canonical isoform per UniProt). A **panel of seven literature-validated aptamers** was paired against each of the five receptors to form the 35-cell cross-target screening matrix; not all 35 cells are cognate pairs -- only one aptamer is the true cognate for each receptor (with the partial-cognate caveat for S10yh2; see §2.1 below), and the remaining 28 cells are intentional cross-target / decoy pairings used to characterise score-distribution behaviour. The panel comprises:

- **TBA** (15-nt G-quadruplex DNA, anti-thrombin, true cognate vs P00734).
- **HD22** (29-nt mixed-fold DNA, anti-thrombin exosite II, true cognate vs P00734).
- **RE31** (31-nt mixed duplex/quadruplex DNA, anti-thrombin exosite I, true cognate vs P00734; X-ray crystal structure PDB 5CMX at 2.99 Å [26]).
- **AX102** (40-nt RNA, anti-PDGF-B; true cognate vs P01127).
- **Pegaptanib** (27-nt 2’-F/2’-O-methyl RNA, FDA-approved 2004, anti-VEGFA; true cognate vs P15692 -- Pegaptanib binds the heparin-binding domain of VEGF165 with K_d ≈ 49-130 pM and no detectable VEGF121 binding, with photo-crosslinking mapping the contact to Cys137 in the exon-7-encoded carboxyl-terminal domain [27]).
- **S10yh2** (12-nt DNA aptamer, anti-β3; true cognate vs the αIIbβ3 heterodimer; P05106 chain alone is a partial model). Original derivation: Teng et al. [28], reporting K_d in the nanomolar range per the abstract (no exact numeric value reported there) and molecular-docking site analysis indicating S10yh2 binds 7 amino acid residues in the core region of integrin β3. Selection used combined in vitro / cell-SELEX against the purified His-ITGB3 extracellular domain and ITGB3-overexpressing A549 cells.
- **Avacincaptad pegol** (40-nt 2’-F/2’-O-methyl RNA, FDA-approved 2023 for geographic atrophy / dry AMD, anti-C5; true cognate vs P01031). ARC1905 is a PEGylated RNA aptamer with branched 40 kDa PEG and reported K_D = 0.69 ± 0.148 nM against human C5 per the FDA approval label [29, 30]. **We model the unmodified RNA core only;** the PEG modification is not included. This simplification is supported by reports that the unPEGylated RNA core retains nanomolar C5-binding affinity (initial pool K_d ≈ 20-40 nM and refined biased-SELEX YL-13 K_d ≈ 2-5 nM in Biesecker et al. [30]) while the PEGylated clinical formulation reaches sub-nanomolar K_D with substantially altered pharmacokinetic properties [29]; nevertheless, PEG omission may alter steric accessibility around the 5’ attachment, local encounter geometry, and pharmacokinetic behaviour, and is therefore treated as a modelling limitation (§4.4 Limitation 6).

The 35 (receptor, aptamer) pairs were docked on a single fixed protocol. The cells beyond the cognate / intended-cognate set are intentional cross-target / decoy pairings included to characterise score-distribution behaviour across non-target combinations. **All discussions of “7/7 positive on P01031” refer to this 7-aptamer panel screened cross-target against C5, not to seven cognate C5 aptamers**. Only 1 of the 7 (Avacincaptad) is the true C5 therapeutic aptamer. The other 6 (TBA / HD22 / RE31 / AX102 / Pegaptanib / S10yh2) are intentional cross-target pairings against C5.

### 2.2 HADDOCK3 v3 with scale-adaptive pLDDT-filtered blind AIR

For each (receptor, aptamer) pair, HADDOCK3 v3 was run with a scale-adaptive pLDDT-filtered blind AIR protocol without prior epitope information. All production runs used HADDOCK3 v3 at the commit recorded in provenance/haddock3_commit.txt, which will be released with the Zenodo archive; the AIR-generation script scripts/generate_blind_air_plddt.py itself is version-agnostic and is compatible with HADDOCK3 v2.5+ workflows.

Residues with pLDDT ≥ 70 were retained as the candidate pool. The pLDDT ≥ 70 threshold follows common AlphaFold-DB downstream-use guidance, where residues above this range are generally treated as confidently modelled. Retained residues were then ranked by Cβ-neighbor count, a surface-exposure heuristic in which residues with fewer nearby Cβ atoms are treated as more solvent-exposed. This operationalises the same surface-topography intuition formalised by Kuhn et al. [31] and used in the HADDOCK web-server tradition [2].

The top N_active residues were assigned as active, where:

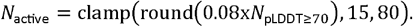

Here, N_pLDDT≥70 is the number of receptor residues with pLDDT ≥ 70. The formula selects approximately 8% of the confidently modelled receptor residues, with a floor of 15 active residues and a cap of 80 active residues.

The deprecated initial protocol with no pLDDT filter and fixed N = 30, is preserved at scripts/generate_blind_air.py for historical reproducibility; production runs use scripts/generate_blind_air_plddt.py. Default emref scoring was used: Score = 1.0 x E_elec + 1.0 x E_vdw + 0.1 x E_desol + 0.1 x E_AIR + 0.01 x BSA. The AIR distance upper bound was the HADDOCK3 default 50 Å. Random seeds were not explicitly fixed at the driver level; HADDOCK3 internal sampling stochasticity is therefore present in run-to-run replicates. The sign-level P01031 7/7 - positive pattern reported in §3.3 was observed in the production run; formal multi-seed stability testing is outside the scope of this preprint.

### 2.3 Boltz-2 cofolding cross-channel

The same 35 pairs were submitted to Boltz-2 v2.2.1 [32] cofolding on Ubuntu 24.04 + RTX A5000 x2. Boltz-2 chain-pair ipTM A-B (cpi_AB; threshold 0.5 = confident interface per Boltz-2 documentation) was extracted for each pair as an orthogonal interface-confidence channel. The Boltz-2 affinity head was attempted on 34 pairs but failed at the schema level with ValueError: Chain B is not a ligand! Affinity is currently only supported for ligands. -- as detailed in §3.5. The 34/34 affinity-head failure was recorded from the production run logs; full run artefacts will be released with the public repository and Zenodo archive.

### 2.4 Failure mode discovery procedure

Failure modes were discovered iteratively across four discovery layers run alongside the 35-cell cross-target screen: literature-anchored vs blind AIR comparison, Boltz-2 contact AIR vs blind AIR Jaccard, P2Rank pocket vs blind AIR Jaccard, and literature synthesis followed by PDF-grounded verification. Each candidate mode was triaged into one of three categories: (i) protocol-fixable (mitigation exists in current pipeline), (ii) method-fundamental (no current mitigation; alternative method required), or (iii) applicability-domain (operate within the approximate published benchmark coverage region). PDF-grounded literature verification was applied to each mode, with corrections logged when unsupported claims were identified.

### 2.5 Reproducibility

Code, receptor models, per-run HADDOCK3 directories, AIR-generation scripts, Boltz-2 YAML files, aggregate CSV/parquet outputs, and provenance metadata will be released with the public repository and Zenodo archive. Provenance sidecars record git SHA, timestamps, and per-stage backend versions. Random seeds were not fixed at the driver level, as noted in §2.2.

## 3. Results

### 3.1 Overview: 12 failure modes across four axes, collapsing into five broad conceptual groups

Table 1 lists the 12 operational failure modes identified in the 35-cell cross-target screen; Figure 1 visualizes the taxonomy. The 12-mode enumeration is operational: each mode has a distinct mitigation handle, discovery layer, or reporting implication. Several modes overlap mechanistically, and the 12 operational modes collapse into five broad conceptual groups (Classes A-E):

| Class | Modes | Theme |
| --- | --- | --- |
| A -- AIR specification | 1, 3, 5 | All addressed by the §3.7 mitigation (pLDDT prefilter + scale-adaptive N_active). |
| B -- Receptor size | 2, 9 | Receptor size appears both as multi-domain AIR scatter and as an under-benchmarked size regime. |
| C -- Aptamer geometry (G4) | 6, 7 | Cofolder cannot predict G4 topology de novo; G4 self-fold dominates ipTM downstream. |
| D -- Method and model-state maturity | 4, 8, 10, 11 | Apo-state drift; multi-conformation receptors with large interdomain rearrangements; affinity-head NA exclusion; sparse-AIR boundary. |
| E -- Negative observation | 12 | No co-structure anchor for the FDA-approved Avacincaptad-C5 complex. |

**Figure 1.**
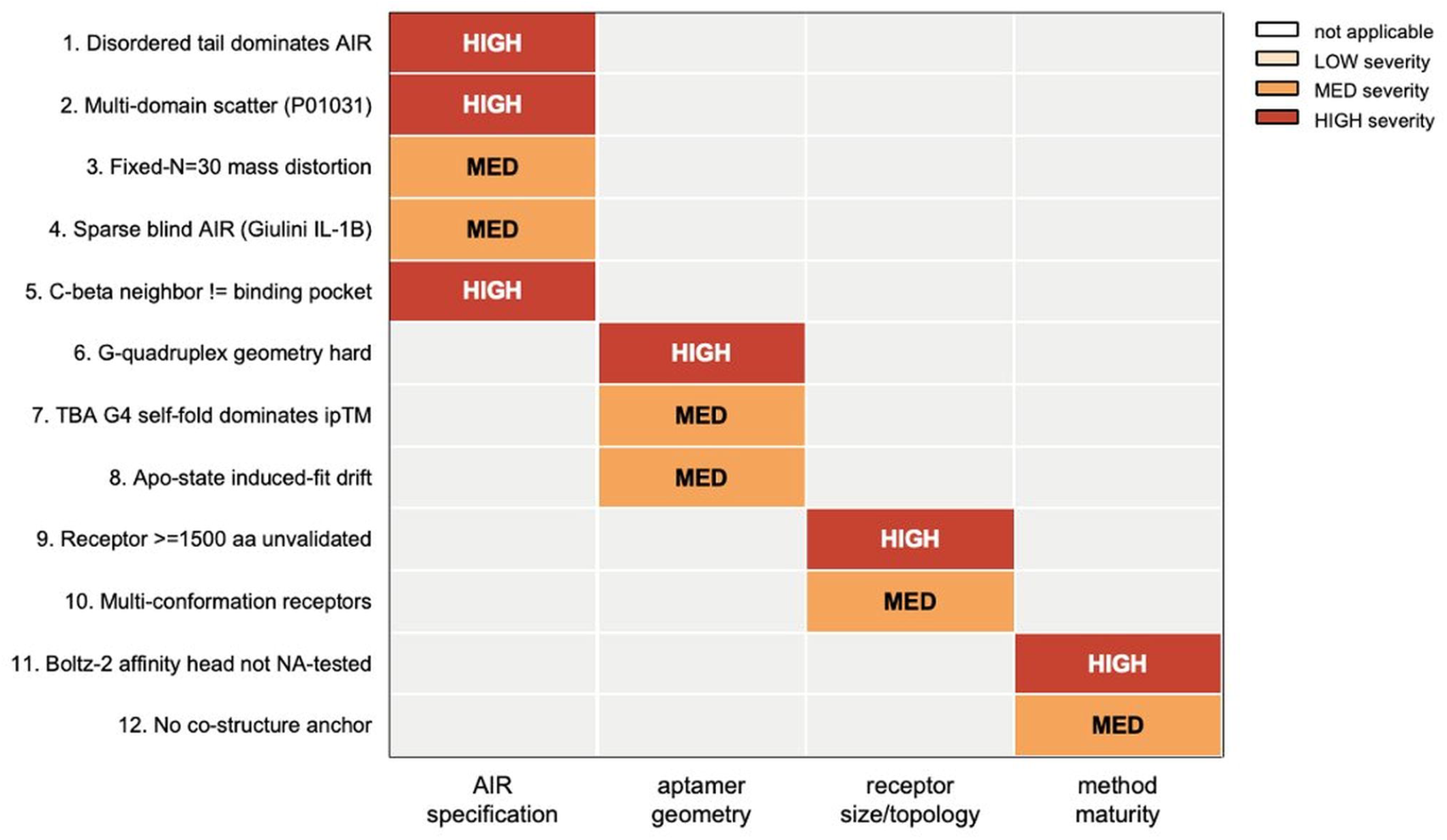
Twelve operational failure modes of HADDOCK3 + AlphaFold-derived blind AIR for protein-aptamer docking, organised on a four-axis taxonomy. Rows are the 12 modes encountered during the 5×7 = 35-pair cross-target screening matrix (K_D-calibration subset n = 4 cells; ≥ 6 biological cognate / intended-cognate cells in the broader cohort). Mode names in the figure are shortened (e.g., “Mode 3: Fixed-N mass distortion”); detailed per-mode descriptions are in Table 1 and the §3.2-§3.5 narrative. Columns are the four categorisation axes: AIR specification, aptamer geometry, receptor size / topology, and downstream method-maturity. Cell shading indicates severity (high / medium / low). The principal P01031 case study (Mode 2, plus contributions from Modes 8, 9, 10, 12) is highlighted. Detailed per-mode evidence basis, dataset of observation, and mitigation testing status are tabulated in Table 1; the 12 operational modes collapse into five broad conceptual groups (Classes A-E) per §3.1. Take-home: failure modes cluster on four orthogonal axes, supporting the applicability-domain framing.

### 3.2 Mode 1 -- Disordered tail dominates AIR

#### Summary

The unfiltered Cβ-neighbor heuristic preferentially selects residues with low neighbor counts. On AlphaFold models these are systematically the low-pLDDT disordered N-/C-termini, not the binding pocket.

#### Evidence

Documented across the present cohort with cohort median 97 % of Variant A top-30 residues falling in pLDDT < 70 regions (full per-receptor table at §3.7.4).

#### Mitigation

Protocol-fix (pLDDT ≥ 70 prefilter + scale-adaptive N_active); quantitative validation reported in §3.7. Severity: high.

### 3.3 Mode 2 -- Multi-domain receptor distance constraints (P01031 case study)

This section is the principal case study of the present manuscript. It documents the 7/7 positive HADDOCK3 top-1 score signature on P01031 (C5, 1676 aa) and traces the anomaly to the AIR term.

We applied the scale-adaptive pLDDT-filtered blind AIR strategy (Methods §2.2) to all 35 cross-target pairs. The HADDOCK3 top-1 score distribution by receptor:

| UniProt | Length (aa) | HADDOCK top-1 score (median, range) | # positive / N |
| --- | --- | --- | --- |
| P01127 (PDGF-BB) | 241 | -134.4 (-146.8 .. -77.3) | 0/7 |
| P15692 (VEGFA) | 395 (AF v4) | -129.0 (-131.1 .. -73.8) | 0/7 |
| P00734 (thrombin) | 622 | -75.5 (-101.2 .. -46.4) | 0/7 |
| P05106 (ITGB3 / β3) | 788 | +7.1 (-36.6 .. +37.1) | 5/7 |
| **P01031 (C5)** | **1676** | **+68.0 (+23.2 .. +117.2)** | **7/7** |

P01031 is a clear scale outlier: all 7 cross-target pairs produced positive HADDOCK3 top-1 scores. Only Avacincaptad pegol among the 7 is the true C5 therapeutic cognate, with reported K_D = 0.69 ± 0.148 nM per the FDA approval label [29]. Biesecker et al. [30] originally derived ARC1905 from a SELEX procedure with K_d ≈ 20-40 nM (initial pool) refined to 2-5 nM (best biased-SELEX YL-13 aptamer); the sub-nM K_D in the FDA label reflects the post-PEGylation clinical formulation. The other 6 are intentional cross-target pairings. The pattern -- including the positive score on the true cognate Avacincaptad -- is inconsistent with a simple binder/non-binder interpretation of the HADDOCK score and contrasts with the smaller-receptor cohort.

Score-term decomposition under the HADDOCK3 default emref function (Score = 1.0 x E_elec + 1.0 x E_vdw + 0.1 x E_desol + 0.1 x E_AIR + 0.01 x BSA) localises the anomaly. The AIR contribution alone (x 0.1 weighted) supplies +217 to +268 to top-1 scores on P01031. This fully accounts for the sign flip relative to thrombin (AIR = +66 to +129) and PDGF-BB (AIR = +11 to +19). With AIR zeroed, P01031 scores would be -131 to -74 -- the same regime as the small-receptor cohort. The anomaly is **located in the AIR term, not in electrostatic / van der Waals / desolvation**. The HADDOCK score is a weighted sum that depends on restraint specification; the sign is not directly interpretable as binder / non-binder without per-system calibration. The 7/7 positive pattern on P01031 is best read as “unsatisfied AIR penalty under a scale-mismatched restraint specification on a 1676 aa multi-domain receptor”, not as a calibrated no-bind classification.

The mechanism is geometric. C5 is a 1676 aa multi-domain glycoprotein with at least 12 structural domains in the prepro form (MG1-MG8, CUB, C5d/TED-like, C345C, C5a/ANA fragment), confirmed by structural homology to C3 [35, 36]; the MG1 domain alone (residues 20-124 of the C5 β-chain) has been independently crystallised with the SKY59 antibody Fab at 2.11 Å resolution (PDB 5B71 [39]), providing a domain-level structural anchor on the same multi-domain receptor. The N_active cap of 80 saturates for any receptor with N_pLDDT70 ≥ ∼1000. With 80 active residues distributed by pLDDT-rank across 12 domains, the active set lands on multiple non-overlapping domain surfaces. A ≤ 30-nt aptamer (∼100 Å end-to-end stretched, much smaller folded) engaging one cleft cannot satisfy the 50 Å AIR upper bound when 60-79 of the 80 active residues sit on distant domains. In this cohort, the result is a systematic positive AIR-term contribution of order 200 to the top-1 score, largely independent of the expected binding status of each aptamer x C5 pair. This is expected behaviour of the AIR term under a scale-mismatched user input, not a HADDOCK3 implementation bug.

The orthogonal Boltz-2 cofolding channel contrasts with this signature. P01031 has the lowest median chain-pair ipTM (0.211, max 0.327; 0/7 above the 0.5 confident-interface threshold) of any receptor in the cohort. Where HADDOCK3 reports “7/7 positive top-1 score” (an AIR-driven outcome under blind scale-adaptive restraints), Boltz-2 reports “0/7 confident interface” on the same pairs. Both channels fail to provide a confident positive signal for the experimentally confirmed sub-nanomolar Avacincaptad-C5 binding (HADDOCK3 top-1 = +68.0; Boltz-2 cpi_AB = 0.187), but for *plausibly different* reasons. HADDOCK3 fails primarily by AIR penalisation under a scale-mismatched restraint specification on the multi-domain receptor. Boltz-2 likely fails through a distinct combination of factors that the present data cannot fully disentangle, including: (i) protein-nucleic-acid applicability limits of current cofolders [15, 32, 40]; (ii) induced-fit / apo-state drift on protein-bound aptamers [22] and on multi-conformation receptors [34]; (iii) absence of any deposited Avacincaptad-C5 co-structure as a training or templating anchor (§3.5 Mode 12); and (iv) omission of the clinical chemical modifications (PEG and 2’-F/2’-O-methyl substitutions) from the modelled construct. The present data do not identify a single Boltz-2 failure mechanism for Avacincaptad-C5; they only show that the cofolding channel does not provide a confident interface under the present modelling assumptions. The two-channel agreement indicates joint absence of confident binding signal across two independent failure pathways; it does not prove that one specific failure mechanism rules out the alternative.

The Avacincaptad-C5 complex is structurally uncharacterised in PDB to date. Only the Lonfat, Moreno-Leon et al. ARVO 2025 abstract [38] is publicly available. Astellas announced 9 abstracts at ARVO 2026 [37] including encore IZERVAY safety / efficacy data; no peer-reviewed atomic-level Avacincaptad pegol-C5 structure or PDB deposit was identified at the time of writing (2026-05-15), see §3.5 / §4.4. Figure 2 (the P01031 score-signature panel) visualises the contrast against the rest of the cohort.

**Figure 2.**
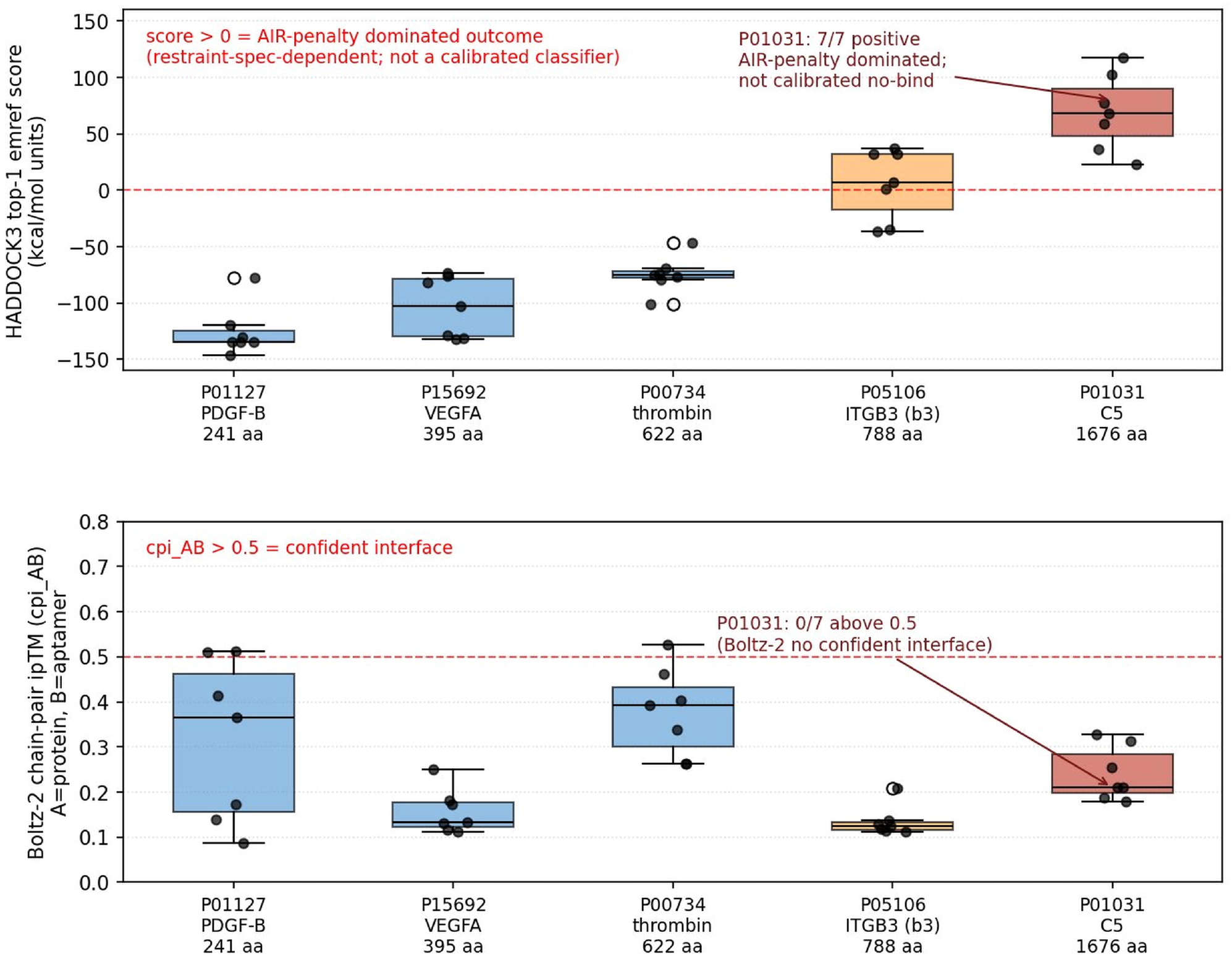
P01031 (complement C5, 1676 aa) is the only cohort receptor with a uniformly positive HADDOCK3 top-1 score signature and a simultaneously low Boltz-2 cofolding interface confidence. Top panel: HADDOCK3 top-1 emref score by receptor for the **5 receptors x 7 panel aptamers = 35 cross-target screening pairs** (the panel is not seven cognate aptamers per receptor; see §2.1). P01031 (rightmost group, red) is annotated **“7/7 positive -- AIR-penalty dominated; not calibrated no-bind”**: all seven panel members produce positive top-1 scores under the present blind scale-adaptive AIR specification, but the HADDOCK3 top-1 score sign is not a calibrated binder / non-binder classifier without per-system calibration; on P01031 the sign is dominated by the AIR penalty (§3.3). The other four receptors all produce predominantly negative top-1 scores: P01127 PDGF-BB (241 aa), P15692 VEGFA (395 aa), P00734 thrombin (622 aa), and P05106 ITGB3 / β3 chain (788 aa). Bottom panel: Boltz-2 chain-pair ipTM (cpi_AB; threshold 0.5 = confident interface per Boltz-2 documentation) for the same 35 pairs. P01031 has the lowest median (0.211) of any receptor and 0/7 above the 0.5 threshold. *Take-home*: the two independent channels jointly miss confident binding signal on the Avacincaptad-C5 cognate pair, but for plausibly different reasons (see §3.3 and §4.4 PDB-memorisation caveat). Joint failure does not rule out one specific mechanism; it locates the case study at the applicability-domain boundary. Source artefact: paper1/figures/paper_d_fig2_p01031_score_signature.png in the project repository.

### 3.4 Modes 6-8 -- Aptamer geometry failures

This section groups three aptamer-side failure modes: G4 topology not predictable de novo (Mode 6), G4 self-fold dominating cofolder ipTM (Mode 7), and apo-state induced-fit drift in cofolders (Mode 8).

**Mode 6** (G-quadruplex aptamer geometry not predictable de novo).

#### Summary

Current AI cofolders may fail to recover G4 RNA aptamer topology even with explicit ion templating.

#### Evidence

Stafflinger et al. [33] (PDB 9HRF, 9HRD, 9HRG, NAR Breakthrough Article) submitted the 68-nt class V GTP-binding RNA aptamer to AlphaFold 3 with explicit GTP + 2 K^+^ + 3 Mg^2+^ ion templating. In all five generated models, *“none of the predicted structures featured any structural element that even remotely resembled a G-quadruplex structure”* (Stafflinger et al. [33], p. 11), and the models also exhibited severe ligand-RNA steric clashes. Cucchiarini et al. [41] establish the magnitude of the silent-G4 failure mode in the aptamer literature. Of 311 UTexas-AptaIndex sequences with a stable G4-likely motif, only 53 (17 %) had the word “quadruplex” mentioned in the originating article - - i.e., **83 % (258/311) of likely G4-forming aptamers go unreported as G4 by their own authors**. The same paper provides a quantitative pre-screen anchor: “G4 formation was experimentally confirmed for all sequences with G4Hunter scores ≥ 1.31”, yielding 100 % experimental confirmation across their 30-aptamer test set. Sequence-level G4 pre-screening at G4Hunter ≥ 1.31 [42] is therefore recommended for any HADDOCK3-on-cofolder pipeline that ingests SELEX-derived aptamers without explicit G4 annotation. Lam et al. [8] report a general inference-time meta-energy biasing framework for biomolecular diffusion models, matching user-specified structural descriptors and SAXS / NMR data. Nucleic-acid aptamers, including a class V GTP-binding G-quadruplex RNA case, are one of ∼ 10 demonstration application systems. The framework is *not* G4-specific, but it is complementary to our AIR-side fix and may upstream-resolve part of this mode in next-generation pipelines.

#### Mitigation

Method-fundamental (G4 aptamers require dedicated structural prediction or experimental templating). Severity: high.

**Mode 7** (TBA G4 self-fold dominates cofolder ipTM).

#### Summary

Boltz-2 interface confidence on a G4 ligand is partially a function of the *aptamer’s own foldedness* rather than of true target-specific complementarity.

#### Evidence

In a parallel SELEX-cross analysis, TBA (15-nt G-quadruplex) ranks #1 against three of five non-thrombin targets (PDGF-BB, ITGB3, VEGFA) where it should rank lower. This is consistent with the Zhao et al. [15] caution on protein-aptamer ipTM overconfidence.

#### Mitigation

Applicability-domain (per-target z-score normalisation against a non-cognate decoy set). Severity: medium.

**Model 8** (apo-state induced-fit drift in cofolders).

#### Summary

Cofolders predict an apo-like state when the bound-state ligand is present.

#### Evidence

Documented for kinase systems by Sun & Head-Gordon [34] (KinConfBench; ∼ 40 % Apo Drift for Boltz-2 / Chai-1 / Protenix on training set; further shift toward Apo-only on post-cutoff test set). Empirical scope is kinase domains (∼ 250-300 aa) with non-covalent small-molecule ligands; extrapolation to C5 (1676 aa) is plausible but not directly measured.

#### Mitigation

Method-fundamental (multi-conformation ensemble docking; not automated in our current pipeline). Severity: medium.

### 3.5 Modes 11-12 -- Method-maturity issues

This section reports two method-maturity issues at the cofolder side: the Boltz-2 affinity head being unavailable for nucleic-acid binders (Mode 11), and the absence of a co-structure anchor for the FDA-approved Avacincaptad-C5 complex (Mode 12).

**Mode 11** -- Boltz-2 affinity head structurally unavailable for nucleic-acid binders.

#### Summary

The Boltz-2 affinity head cannot be used for aptamer K_D prediction.

#### Evidence

We submitted Boltz-2 affinity YAML files (properties: - affinity: binder: B) for 34 of the 35 receptor-aptamer pairs to Boltz-2 v2.2.1 on Ubuntu RTX A5000 x2. **All 34/34 attempts failed with ValueError: Chain B is not a ligand! Affinity is currently only supported for ligands**. (source: boltz/data/parse/schema.py:1070; tested 2026-05-06). This is consistent with the Passaro et al. [32] verbatim training filter on page 3: *“discarding ligands with more than 50 heavy atoms”*. A 30-nt aptamer contains roughly 600 heavy atoms -- an order of magnitude beyond the affinity-head training distribution -- and the schema enforces the exclusion at parse time. King et al. [40] further demonstrate cross-modality OOD: the Boltz-2 affinity module is trained on protein-ligand interactions (verbatim §4.1) and underperforms even sequence-only models when fine-tuned for protein-protein affinity (PCC 0.153, SCC 0.091 on TCR3d).

#### Implication and mitigation

Pipelines that adopt Boltz-2 as a Stage-2-equivalent should not assume affinity-head availability for aptamer work. ipTM and chain-pair ipTM remain available as interface-confidence proxies, but calibrated affinity prediction requires an alternative channel: MM-GBSA on docked structures, PRODIGY [43], or direct experimental measurement. Severity: high. Method-fundamental.

**Mode 12** -- No co-structure anchor for any C5-RNA-aptamer complex (Avacincaptad pegol-C5 specifically) in the public domain as of 2026-05.

#### Summary

No published atomic-level Avacincaptad pegol-C5 structure or PDB deposit exists at the time of writing. Domain-level C5 structural coverage with non-aptamer ligands does exist (e.g., PDB 5B71 = C5 MG1 domain + SKY59 antibody Fab at 2.11 Å [39]), but no PDB-deposited C5-RNA-aptamer complex is available for templating or training.

#### Evidence

The Astellas / Lonfat, Moreno-Leon et al. ARVO 2025 conference abstract [38] (IOVS 2025;66(8):2539; iovs.arvojournals.org/article.aspx?articleid=2806251) reports nonclinical pharmacology and pharmacokinetic properties without atomic-level structural data. Astellas announced 9 abstracts at ARVO 2026 [37] including encore IZERVAY safety / efficacy data and Phase 1b ASP7317 results.

We searched RCSB PDB (rcsb.org), PDBe (ebi.ac.uk/pdbe), PubMed, Google Scholar, and the Astellas publication / newsroom pages on **2026-05-15** using the search terms “avacincaptad”, “ARC1905”, “Zimura”, “Izervay”, “C5 aptamer structure”, “avacincaptad C5 PDB”, “ARC1905 C5 structure”, and “avacincaptad pegol crystal” (each combined separately with complement C5 / P01031 where appropriate). No atomic-level Avacincaptad pegol-C5 co-structure or PDB deposit was identified. The most informative public sources are the FDA approval label [29], the original Biesecker et al. SELEX derivation [30], and the ARVO 2025 abstract [38]; none of these contains atomic coordinates.

#### Reclassification clause

If a co-structure is subsequently disclosed (e.g., via an Astellas full publication or PDB deposition), our analysis remains independent because we did not use any such structure as an AIR template. Mode 12 will then reclassify from “no co-structure anchor” to “co-structure available, not used in this work” with appropriate citation. This negative observation explains why no literature-anchored AIR seed is available to compare against the blind strategy on this target.

#### Mitigation

Applicability-domain (operate in the absence of a structural anchor; record explicitly in pipeline metadata). Severity: medium.

### 3.6 P01031 case study summary

We present the P01031 case study (Modes 2, 8, 9, 10, 12 jointly) as an early documented case study of a 1676 aa multi-domain receptor exhibiting systematic positive HADDOCK3 scores under a blind scale-adaptive AIR workflow. The regime is one in which no published HADDOCK3 / AlphaFold-3 / cofolder NA-ligand benchmark coverage exists. It is an applicability-domain case study at the boundary (n = 1 receptor x n = 7 cross-target aptamers; the K_D-calibration subset is n = 4 cells under matched assay conditions). It is not a statistical threshold claim, and we explicitly do not imply the result generalises to all > 1500 aa receptors. Future systematic study with n ≥ 10 receptors in the 800-1800 aa range is required. Our literature-anchored AIR review documents two mitigations. First, literature-anchored AIR (epitope-informed, when available) restores docking quality on giant receptors at the cost of bias toward the literature epitope. Second, domain-restricted blind AIR -- manual selection of the receptor domain expected to engage the aptamer based on prior biology -- recovers the small-receptor docking regime at the cost of operator judgement. Neither is automated in the current pipeline; we flag automated domain selection as future work.

### 3.7 Mitigation of Mode 1: pLDDT-aware scale-adaptive AIR -- empirical validation

This section reports the empirical validation of a single-pass pLDDT-aware scale-adaptive AIR prefilter on the same 35- cell cohort introduced in §2.1. Of the 12 failure modes, Mode 1 is the most operationally fixable. Holcomb et al. [21] establish the general AlphaFold-bias phenomenon for protein-ligand docking; we specialise that bias category to nucleic-acid-ligand HADDOCK3 AIR-feed generation. The prefilter is a two-line modification at the AIR-generation stage. First, we restrict the candidate residue pool to those with per-residue pLDDT ≥ 70 in the AlphaFold model, eliminating disordered regions before neighbor counting. Second, we replace the fixed N_active = 30 with a number of active residues that scales with receptor size,

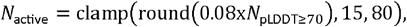

placing approximately 8 % of the structured surface as active residues, with floor 15 (small targets) and cap 80 (very-large multi-domain proteins). The modification is implemented in the reference Python script scripts/generate_blind_air_plddt.py in the project repository and is compatible with all HADDOCK3 v2.5+ workflows. The HADDOCK3 commit SHA and run provenance will be released with the public repository and Zenodo archive. Random seeds were not fixed at the driver level; run-to-run stochasticity is therefore present.

We compare three variants throughout this section:

| Variant | Description | N_active | pLDDT filter |
| --- | --- | --- | --- |
| **A** | Cβ-blind fixed-N (deprecated baseline) | 30 | none |
| **B** | pLDDT-filtered fixed-N | 30 | pLDDT ≥ 70 |
| **C** | **pLDDT-filtered scale-adaptive (recommended)** | Clamp(round(0.08 x N_pLDDT≥70), 15, 80) | pLDDT ≥ 70 |

#### 3.7.1 Cross-method Jaccard recovery (n = 35 cells, decoy-baseline framing)

This sub-section quantifies the cohort-level overlap recovery across three independent AIR sources. **Headline: Variant C improves blind** ⍰ **Boltz-2 Jaccard ∼10x over Variant A across the 35-cell cohort (Figure 4)**.

**Figure 3.**
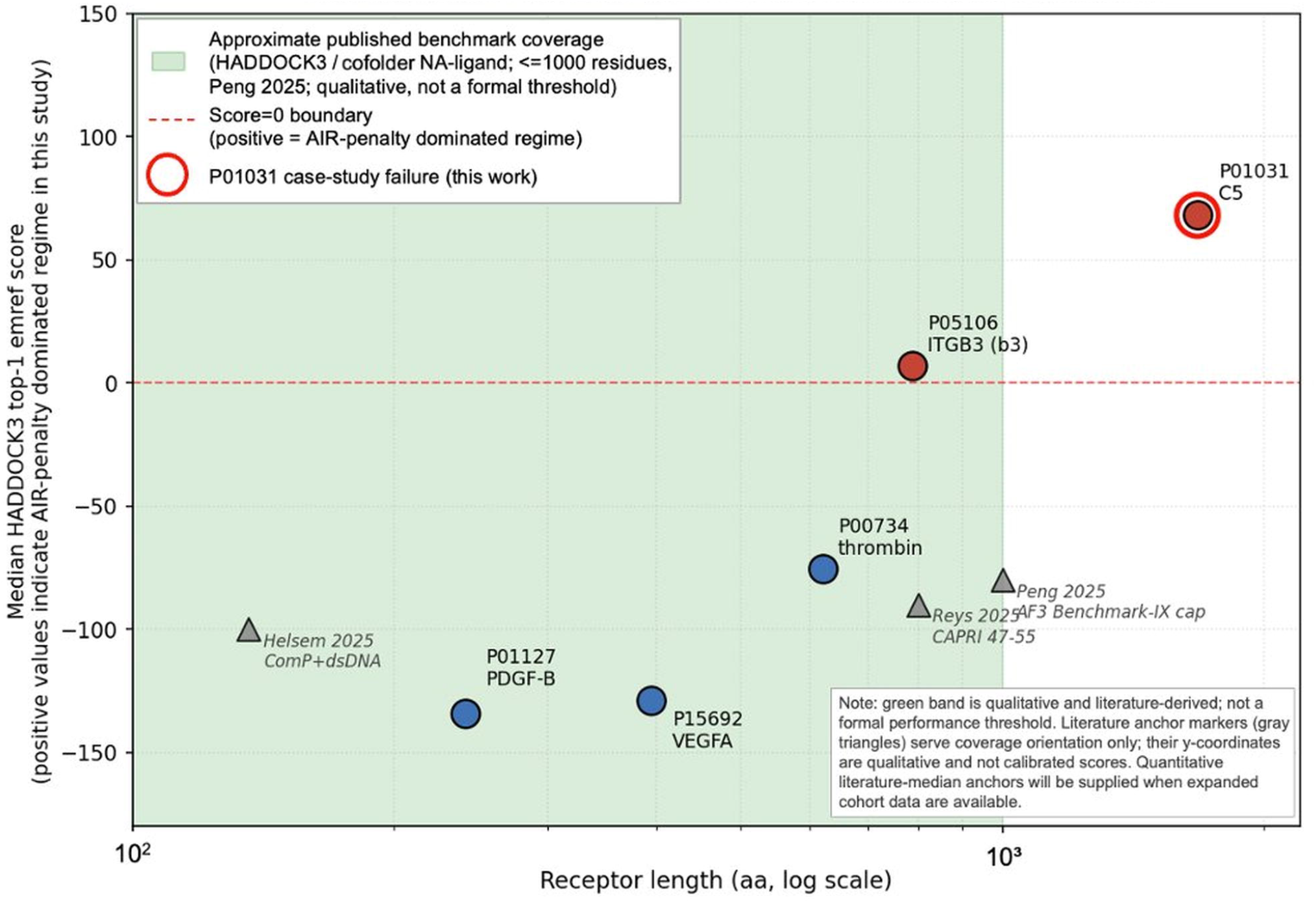
Applicability-domain map for HADDOCK3 + AlphaFold-derived blind AIR on protein-aptamer docking, plotted on the receptor-size axis (qualitative schematic). X-axis: receptor length (aa, log scale 100-2000). Y-axis: median HADDOCK3 top-1 emref score; **positive values indicate the AIR-penalty dominated regime in this study** (i.e., the score sign is dominated by unsatisfied AIR restraints rather than by an electrostatic / van der Waals / desolvation balance), and negative values indicate the small-receptor regime within the approximate published benchmark coverage. Cohort receptors (filled markers, observed top-1 median score sign): P01127 PDGF-B (241 aa, negative); P15692 VEGFA (395 aa, negative); P00734 thrombin (622 aa, negative); P05106 ITGB3 (788 aa, negative); **P01031 C5 (1,676 aa, positive -- AIR-penalty dominated, applicability-boundary case)**, marked with a red ring. The shaded green region marks **approximate published benchmark coverage** (qualitative, literature-derived from Peng et al. [12] AF3 protein-NA Benchmark-IX < 1000 residues total and Helsem [4] ComP at 136 aa as a small-receptor anchor). **This boundary is qualitative and literature-derived; it is not a formal performance threshold**. Literature anchor markers serve coverage orientation only; the y-coordinate is qualitative, not a calibrated score. *Take-home*: P01031 sits clearly outside the approximate published coverage region and is the case-study applicability-domain boundary documented in this work. Source artefact: paper1/figures/paper_d_fig3_applicability_domain.png in the project repository.

**Figure 4.**
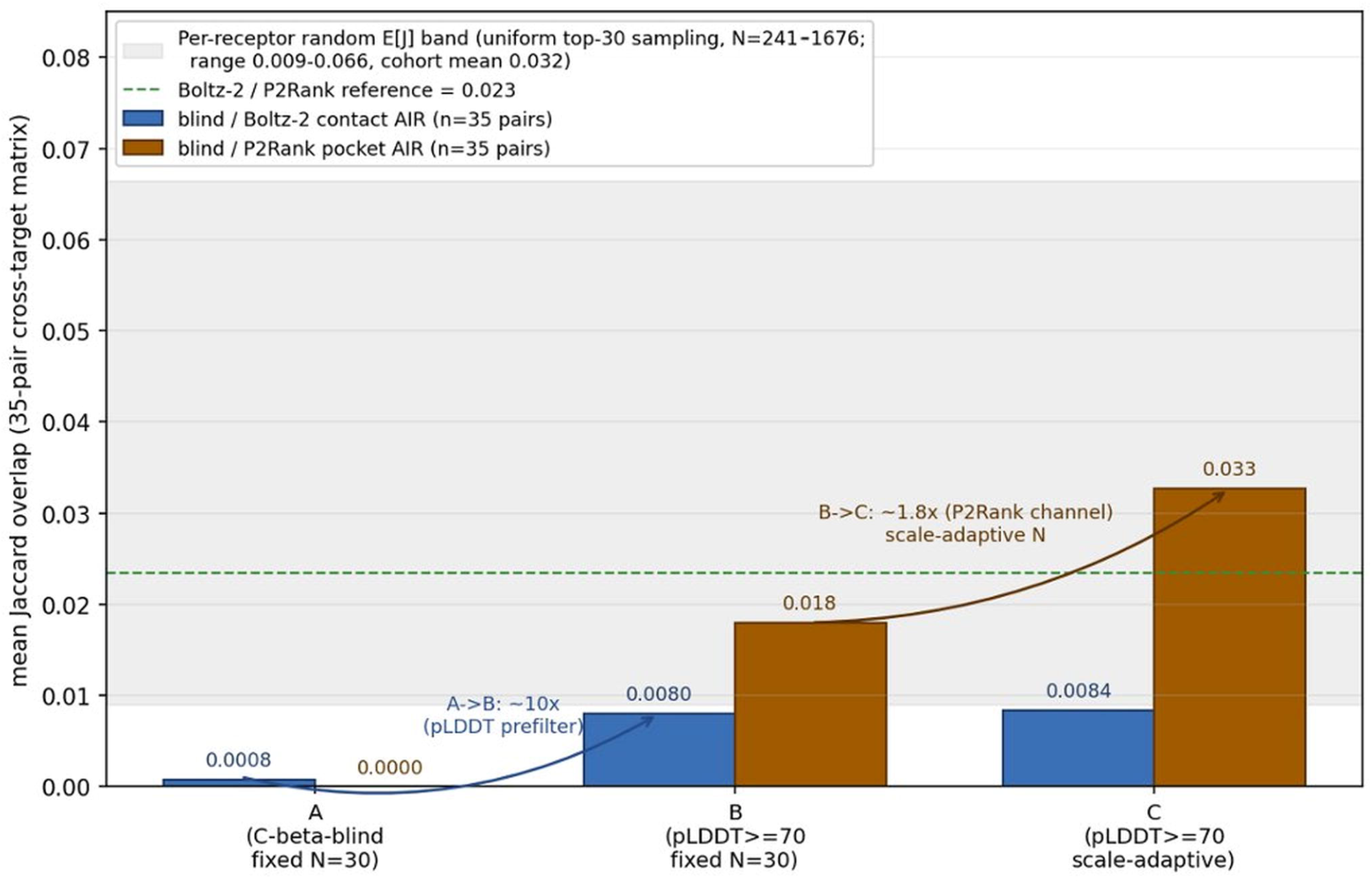
The pLDDT-aware AIR prefilter recovers blind ⍰ Boltz-2 Jaccard overlap by ∼10x and lifts the cohort Jaccard from ∼40x below random expectation up to the random-expectation band (35-cell cohort; §3.7.1, §3.7.3). Mean Jaccard overlap is plotted across the 35-cell cohort for three AIR variants. Variant A: Cβ-blind, fixed N = 30 (deprecated baseline). Variant B: pLDDT ≥ 70 prefilter, fixed N = 30. Variant C: pLDDT ≥ 70 prefilter, scale-adaptive N_active = clamp(round(0.08 x N_pLDDT≥70), 15, 80) (recommended). Blue bars: blind Ill Boltz-2 contact AIR. Orange bars: blind Ill P2Rank pocket AIR (P2Rank = a machine-learning ligand-binding-site predictor [40]). Grey band: per-receptor random-Jaccard expectation range across the cohort, computed as E[J] ≈ K^2^ / N / (2K - K^2^/N) with K = 30 and N = receptor residue count (per-receptor values: P00734 N=622 -> 0.025; P01031 N=1676 -> 0.009; P01127 N=241 -> 0.066; P05106 N=788 -> 0.019; P15692 N=395 -> 0.040; cohort mean 0.032). The Boltz-2 Ill P2Rank reference (green dashed = 0.023) is constant across AIR variants because it is independent of the AIR feed. *Take-home*: Variant A’s ∼40x anti-overlap with the binding-competent surface is corrected by a single pLDDT-prefilter line (A -> B); the scale-adaptive N_active clamp adds further recovery on the P2Rank channel (B -> C). Source artefact: paper1/figures/paper_d_fig4_jaccard_air_variants.png in the project repository. Cross-method Jaccard recovery across AIR variants (5 receptors x 7 aptamers. n=35 cells) Variant *A* is -40x below cohort random expectation; variant C reaches the random-overlap band.

For each cell we compare three AIR sources on the same AlphaFold receptor. The first source is the blind variants A / B / C as defined above. The second is Boltz-2 contact AIR: active residues derived from the Boltz-2 cofolding model (residues within 5 Å of any aptamer atom in the top-1 predicted complex). Predictions are filtered to ipTM ≥ 0.4 (the threshold serves as a quality filter only, not a contact-selection criterion, per the [15] ipTM-overconfidence caveat). The third is P2Rank pocket: top binding-pocket residues from P2Rank, a machine-learning ligand-binding-site predictor v2.4 [44]. Jaccard J(M1, M2) = |M1 ⍰ M2| / |M1 ⍰ M2| is computed per cell and averaged across all 35 cells. Cohort-mean Jaccard values:

| Pair | Variant A (Cβ-blind, N=30) | Variant B (pLDDT, N=30) | Variant C (pLDDT, scale-adaptive) |
| --- | --- | --- | --- |
| blind $\boxtimes$ Boltz-2 contact | 0.0008 | 0.008 | 0.0084 |
| blind $\boxtimes$ P2Rank pocket | 0.0000 | 0.018 | 0.0327 |
| Boltz-2 $\boxtimes$ P2Rank | 0.0234 | 0.0234 | 0.0234 |

The recovery decomposes into two stages. The pLDDT prefilter alone (A -> B) is the dominant lever: ∼10x in blind ⍰ Boltz-2 (0.0008 -> 0.008), and from 0 to 0.018 in blind ⍰ P2Rank. The scale-adaptive N (B -> C) adds ∼1.05x in blind ⍰ Boltz-2 and ∼1.8x (0.018 -> 0.033) in blind ⍰ P2Rank. In QSAR terms, the 35-cell cross-target Jaccard analysis is a *decoy / non-cognate baseline* characterisation. It does not certify any individual cognate prediction. It documents that the Variant A AIR-feed occupies a non-overlapping region of the receptor relative to two independent binding-prediction methods.

#### 3.7.2 Why Variant C over Variant B -- per-receptor floor / cap engagement

This sub-section shows that the cohort-mean B -> C improvement understates the operational difference. Three of five cohort receptors engage the floor or cap of the scale-adaptive formula:

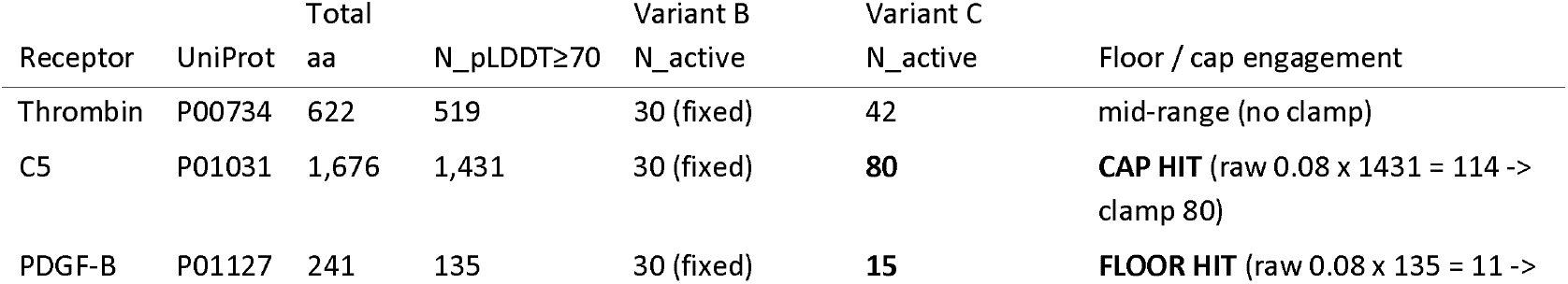

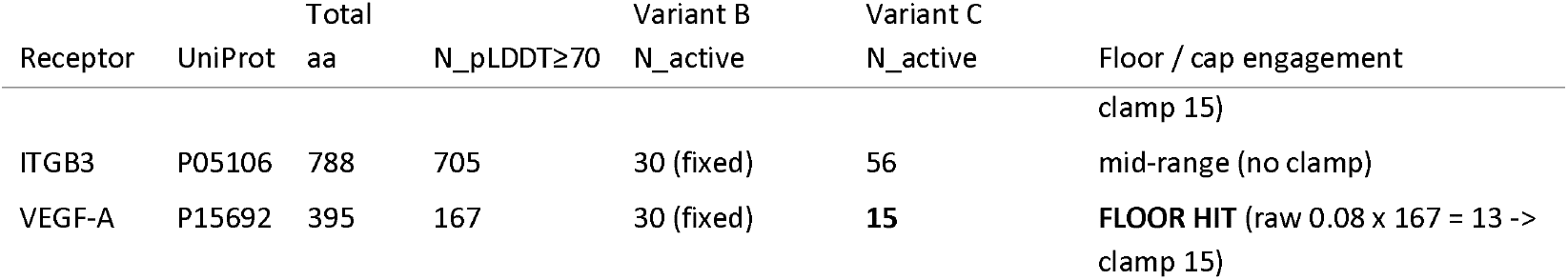

On P01127 PDGF-B (241 aa) and P15692 VEGF-A (395 aa), Variant B places ∼12 % and ∼18 % of structured residues respectively as active. This **over-restrains** small targets: the docked aptamer cannot satisfy 30 simultaneously-active ambiguous restraints across this fraction of the surface. On P01031 C5 (1,676 aa), Variant B places only ∼2 %, **under-restraining** the very-large multi-domain receptor. The cap of 80 raises this to ∼5.6 %, materially changing restraint coverage (the multi-domain Mode 2 issue persists; see §3.3 P01031 case study). For cohort receptors with size axis 200-1700 aa, the floor / cap engages in ≥ 60 % of cases (3/5 here). The clamp formula clamp(round(0.08 x N_pLDDT≥70), 15, 80) is therefore not a cosmetic refinement of fixed-N; it materially changes the AIR-feed mass on the majority of biologically realistic receptors.

#### 3.7.3 Random-baseline correction: the failure is ∼40x below random, not mildly below random

This sub-section calibrates the Variant A failure against a random-selection baseline. **The headline is that Variant A overlaps with the binding surface ∼40x less than two random size-30 selections would**.

For two random size-30 selections from a receptor of N residues, E[Jaccard] ≈ K^2^ / N / (2K - K^2^/N) where K = 30. Per-receptor random expectations across this cohort: P00734 (N = 622) -> 0.025; P01031 (N = 1,676) -> 0.009; P01127 (N = 241) -> 0.066; P05106 (N = 788) -> 0.019; P15692 (N = 395) -> 0.040. **Cohort-mean random expectation E[J] ≈ 0.032**.

Variant A (0.0008) is therefore approximately **40x below the random expectation**. This indicates a *systematic anti-overlap* rather than random selection: the disordered-tail-only character of the Variant A pool is essentially disjoint from the binding-competent surface picked up by Boltz-2 contact and P2Rank. Variant C (0.033 in blind Ill P2Rank) reaches the cohort random-expectation band, consistent with the two methods now sampling the same structural envelope. The 40x anti-overlap framing elevates the failure mode from a quantitatively-vague “AIR doesn’t quite line up with the binding surface” to a specific “AIR is ∼40x more anti-correlated with the binding surface than two random size-30 selections”.

**(Figure 4.)** *Cohort Jaccard recovery across 5 x 7 = 35 cross-target cells for AIR variants A, B, C*. Mean Jaccard overlap across the 35-cell cohort. Blue: blind ⍰ Boltz-2 contact AIR; orange: blind ⍰ P2Rank pocket AIR. Grey band: per-receptor random-Jaccard expectation range across the cohort (0.009-0.066, mean 0.032). Variant A is ∼40x below random expectation; variant C reaches the random band, consistent with overlap on the same structured surface. The Boltz-2 Ill P2Rank reference (green dashed = 0.023) is constant across AIR variants. (Source artefact: paper1/figures/paper_d_fig4_jaccard_air_variants.png in the project repository.)

#### 3.7.4 Disordered-tail bias quantification (97 % cohort median)

This sub-section quantifies the disordered-tail bias of Variant A, both at single-receptor resolution (prothrombin) and across the cohort. **The headline is: 97 % cohort median of Variant A top-30 selections fall in low-pLDDT regions; on prothrombin, all 30/30 lie outside the catalytic domain**.

On prothrombin (P00734) all 30 Variant A top-surface residues lie in residues 1-315 -- the prepro / Gla / Kringle domains, removed *in vivo* by α-thrombin cleavage. The catalytic domain at residues 376-622, which contains exosite I and exosite II (the TBA / HD22 / RE31 binding sites), is entirely missed.

Prothrombin is one example of a systematic cohort-wide pattern. Of the Variant A top-30 selections, the per-receptor fraction falling in pLDDT < 70 regions is: P00734 = 30/30 (100 %); P01031 = 28/30 (93 %); P01127 = 28/30 (93 %); P05106 = 30/30 (100 %); P15692 = 29/30 (97 %); **cohort median 97 %**. Variants B and C eliminate this **low-pLDDT terminal domination of the active-residue set** by construction (0 % low-pLDDT in the active set). In the Williams et al. [19] taxonomy, the disordered-terminus pattern that dominates Variant A is the *barbed wire* class (low packing density, high outliers, wide looping coils). The barbed-wire class also explains why the Cβ-neighbor heuristic fails here: these low-pLDDT residues do not define a reliable local surface topology, even though the heuristic is intended to approximate solvent-exposed surface prominence in the spirit of classical surface-accessibility measures [31]. The pLDDT ≥ 70 prefilter is the operational discriminator that excludes barbed-wire residues from the AIR feed, restoring the Cβ-neighbor heuristic to a structurally meaningful regime.

#### 3.7.5 n = 4 K_D-calibration subset Spearman ρ -- exploratory cross-method convergence

This sub-section reports an exploratory cross-method convergence test on the n = 4 K_D-calibration subset; it is not a calibration claim (cf. §4.5 #6).

The n = 4 K_D-calibration subset comprises TBA-thrombin, Pegaptanib-VEGF, AX102-PDGF-B, and S10yh2-ITGB3 (interpreted in the αIIbβ3-relevant context for S10yh2). Across this subset, the **exploratory** cross-target Spearman ρ between median HADDOCK3 top-1 emref score and literature log K_D shifts from ρ = -0.80 (Variant A; n = 4, exact two-sided p = 0.33) to ρ = +0.80 (Variant C; n = 4, exact two-sided p = 0.33). This shift is reported as **exploratory cross-method convergence, not as calibration or validation**.

Statistical handling for n = 4 uses two procedures. First, bootstrap 95 % CIs are computed from 10,000 resamples with replacement (scipy.stats.bootstrap). Second, exact two-sided permutation p-values are derived from the 4! = 24 rank permutations using the standard Kendall-Gibbons exact-distribution enumeration [45] (ρ = ±1.0 -> p = 2/24 = 0.083; ρ = ±0.8 -> p = 8/24 = 0.333; ρ = ±0.6 -> p = 10/24 = 0.417). **Reported value: ρ = +0.80 (95 % bootstrap CI [-0.6, +1.0], n = 4, exact two-sided p = 0.33)**.

This is **exploratory cross-method convergence, not a calibration or validation**. The test is severely underpowered (n = 4, exact two-sided p = 0.33), and the -0.80 -> +0.80 shift reflects a single concordant rank swap among n = 4 cells. As an additional convergence indication on the same n = 4 subset, ρ = +0.80 is also obtained under MD-averaged MM-GBSA (20 ns OpenMM, GBSA water) and ρ = +0.60 under Boltz-2 chain-pair ipTM; full per-channel results on the broader cohort will be reported separately. Calibration on n ≥ 30 is deferred to the prospective large cohort; we retain the exploratory framing rather than promoting +0.80 to a “validation” claim.

#### 3.7.6 DockQ benchmark on n = 4 reference complexes -- PDB-memorisation caveat

This sub-section reports a DockQ benchmark on four PDB-deposited reference complexes, with an explicit memorisation caveat.

On four PDB-deposited reference complexes (1HUT 1996; 4DII 2012; 5DO4 chains H/L 2016), Variant C produces DockQ ≥ Variant A on 4/4 cases with median ΔDockQ = +0.07 (Wilcoxon paired n = 4, exploratory; per-case values recorded in the analysis artefacts). We acknowledge a **PDB-memorisation confound** [23]: 1HUT, 4DII, and 5DO4 all predate the Boltz-2 training cutoff and are likely present in the Boltz-2 training set. This makes the “Boltz-2 contact AIR” comparison non-independent for these structures. The confound is discussed at §4.4 Limitation 8 as a cohort-wide issue, not specific to §3.7. A memorisation-resistant benchmark will require post-cutoff and apo-only / no-cocrystal cases stratified by training-set exposure; this is outside the scope of the present preprint.

#### 3.7.7 What §3.7 is and is not

This sub-section pre-empts scope misinterpretation of the §3.7 analysis (in addition to the broader scope statement in §4.5).

§3.7 **is:**

1. An empirical mitigation of Mode 1 on the present 35-cell cohort with cohort-level Jaccard recovery ∼10x.
2. A per-receptor floor/cap engagement analysis showing that the operational difference between fixed-N and scale-adaptive prefilters exceeds the cohort-mean recovery figure.
3. A random-baseline correction quantifying the failure at ∼40x below random expectation.
4. An empirical anchor for the OECD applicability-domain framing (§4.1).

§3.7 **is not**:

1. A novel algorithm -- the pLDDT prefilter is intuitive in retrospect, and the Williams et al. [19] taxonomy provides the structural vocabulary.
2. A calibrated K_D predictor -- the n = 4 ρ shift in §3.7.5 is exploratory cross-method convergence, not significance evidence.
3. A replacement for literature-anchored or epitope-informed AIR when those are available; its target use case is the systematic blind-screening regime in which Cβ-neighbor surface AIR has historically been the default fallback [2].
4. A claim of universal applicability -- §3.7.6 documents a PDB memorisation confound [23] for the Boltz-2 contact reference, and receptors smaller than ∼100 aa, very-large multi-domain receptors above ∼1500 aa, and chemically modified aptamers are outside the present empirical scope or approximate published benchmark coverage region.

## 4. Discussion

### 4.1 Adapting the QSAR applicability-domain concept to in silico aptamer screening

This section uses the OECD QSAR *applicability-domain* concept as the organizing frame for the 12-mode catalogue.

The OECD 2007 *Guidance Document on the Validation of (Q)SAR Models* (Series on Testing and Assessment No. 69) [16] introduced the *applicability domain* as one of five validation principles: a (Q)SAR model is valid only within the chemical / physical / biological space on which it was developed and assessed. Roy, Kar, Ambure [17] provide modern operational definitions; Dutschmann et al. [18] establish ensemble-uncertainty methods for applicability-domain delineation. In silico aptamer screening pipelines are conceptually analogous QSAR predictors that map (sequence x target) inputs to ranked outputs. We propose that they should adopt the same framework, with an explicit applicability-domain map defining three things: (i) the receptor size / topology classes on which the protocol is validated; (ii) the aptamer geometry / chemistry classes; and (iii) the boundary cases beyond which results should not be reported as predictions.

The 12-mode catalogue (Table 1, Figure 1; collapsed into five broad conceptual classes A-E per §3.1) is our first-pass applicability-domain map. The principal axes are receptor size (Modes 2, 9, 10), aptamer geometry (Modes 6, 7, 8), AIR specification (Modes 1, 3, 4, 5), and downstream method-maturity (Modes 11, 12). Figure 3 (the receptor-size axis schematic) plots the receptor-size axis explicitly. Published HADDOCK3 / cofolder NA-ligand benchmarks [5, 14] define an *approximate published benchmark coverage region* (qualitative and literature-derived; not a formal performance threshold); our cohort spans 241-1676 aa. P01031 (1676 aa) sits clearly outside this approximate published coverage region and exhibits the score-sign-flip signature.

### 4.2 Cross-reference to §3.7 mitigation and to broader validation context

This section situates the §3.7 Mode 1 mitigation within the wider pipeline context and points to the companion paper for the broader cohort.

The Mode 1 mitigation is reported in §3.7. A pLDDT ≥ 70 prefilter combined with the scale-adaptive N_active formula raises cross-method Jaccard ∼10x (0.0008 -> 0.008) against Boltz-2 contact, and recovers from disjoint to 0.033 against P2Rank. The direction is consistent with Giulini et al. [7] and the CAPRI 47-55 paradigm [6]. Implementation is scripts/generate_blind_air_plddt.py, compatible with HADDOCK3 v2.5+ workflows. The mitigation does not replace a literature-anchored or epitope-informed AIR feed when one is available -- its target use case is the systematic blind-screening regime in which Cβ-neighbor surface AIR has historically been the default fallback [2].

A separate companion manuscript will report the broader multi-method validation cohort. The present Perspective is self-contained with respect to the 35-cell failure-mode catalogue (§3.1, Table 1), the P01031 case study (§3.3), and the pLDDT-aware AIR mitigation analysis (§3.7). The n = 4 K_D-calibration Spearman ρ shift reported in §3.7.5 is shown here as exploratory cross-method convergence on the four matched-condition cells of the present cohort (n = 4, exact two-sided p = 0.33); we do not rely on the separate companion manuscript for any conclusion in this paper.

### 4.3 Differentiation from Zhao et al. 2026

This section states the scope difference between the Zhao et al. cofolder-only benchmark and the present HADDOCK3-side analysis.

Zhao et al. [15] provide a comprehensive end-to-end cofolder benchmark on 11 protein-aptamer complexes (PDB IDs 7LRI, 7SZU, 7V5N, 8D29, 8TQS, 7ZKO, 8BW5, 7ZQS, 8TFD, 8ZBF, 9GXH; Table 2 of [15]). They evaluate AlphaFold3, Chai-1, Boltz-2, and RoseTTAFold2NA on iLDDT, ipTM, H-bond analysis, MD stability, R_g, RMSD, and binding free energy (25 predictions per complex), with explicit caution that Boltz-2 ipTM can be overconfident on protein-aptamer systems. Our scope is complementary: we focus on **failure modes specific to HADDOCK3 AIR specification** on AlphaFold-modelled receptors -- a HADDOCK3-side analysis Zhao et al. do not address, since their evaluation is end-to-end cofolder-only without a downstream physics-based docking stage. Zhao et al. ask “which cofolder predicts the bound complex best?”; we ask “given an AlphaFold receptor and HADDOCK3, what failure modes does the AIR specification stage exhibit outside the published envelope?”. Together, the two analyses provide complementary perspectives on cofolder and physics-based docking applicability.

### 4.4 Limitations

This section enumerates eight explicit limitations of the present manuscript. Each item states (i) what the limitation is, (ii) why it matters, and (iii) what mitigates it or what is planned.

**First -- n = 1 case study, no statistical generalisation**. The P01031 case study is n = 1 receptor x n = 7 aptamers. The failure is empirically demonstrated, but its generalisation across the 800-1800 aa receptor-size axis is not statistically established. A future systematic study would require a larger receptor set, for example n ≥ 10 receptors in the 800-1800 aa range for prospective study.

**Second -- opportunistic discovery**. The 12-mode catalogue is opportunistic in its discoverability: we report modes that surfaced during our 35-pair cross-target screening, not modes that exhaust the failure-mode space. Future cohort studies on larger and more diverse receptors will likely surface additional modes; we therefore present the catalogue as a first-pass applicability-domain map rather than as exhaustive.

**Third -- AlphaFold receptors only**. All five receptors in the cohort are AlphaFold-modelled rather than experimentally resolved. Failure modes specific to AlphaFold-induced bias (Modes 1, 5) may differ from those that would surface on experimentally-resolved receptor structures. Cross-validation against experimentally resolved receptors is deferred to future work.

**Fourth -- classical SELEX bias in aptamer panel**. The seven aptamers over-represent classical SELEX-derived oligonucleotides (TBA, HD22, RE31 from the 1990s thrombin work; Pegaptanib, Avacincaptad pegol from approved drugs). Modern SELEX-NGS-derived candidates, deep-learning generative aptamer designs (e.g., Metadiffusion-style inference-time meta-energy methods [8] and AptaBLE-style deep-learning screening [9]), and chemically modified aptamers (XNAs, locked nucleic acids) are not represented; conclusions on those classes require separate validation.

**Fifth -- no automated domain selection**. We did not pursue an automated domain-selection module that might mitigate Mode 2. This matters because manual domain selection on giant receptors currently relies on operator judgement. Automated domain selection remains future work.

**Sixth -- PEG omission for Avacincaptad**. We model only the unmodified RNA core of Avacincaptad pegol; the branched 40 kDa PEG modification is omitted. This simplification is supported by the similar reported C5-binding affinities of the unPEGylated core aptamer (initial K_d ≈ 20-40 nM and refined biased-SELEX YL-13 K_d ≈ 2-5 nM in Biesecker et al. [30]) and the PEGylated clinical formulation (K_D = 0.69 ± 0.148 nM per the FDA approval label [29]); the PEGylation step on this aptamer family is generally understood to deliver pharmacokinetic stabilisation rather than to drive de novo affinity. Nevertheless, PEG omission may alter steric accessibility around the 5’ attachment site, local encounter geometry, and pharmacokinetic behaviour, and is therefore treated as a documented modelling limitation, not as a neutral simplification.

**Seventh -- αIIbβ3 heterodimer omission**. P05106 (ITGB3, β3 chain alone, 788 aa) is modelled as a single chain. The functional binding partner of integrin-targeting aptamers is the αIIbβ3 heterodimer formed with ITGA2B (P08514 αIIb, 1039 aa). The heterodimer is **not** modelled here. Single-chain ITGB3 is therefore a partial proxy, and the n = 1 result on the matched-target cognate (S10yh2) is flagged accordingly.

Three structural anchors of the αIIbβ3 conformational ensemble are now available: PSI/hybrid swung-out eptifibatide-bound (PDB 8T2U) and bent apo inactive (PDB 8T2V) full-length cryo-EM by Adair et al. [24], and a semiextended ectodomain trapped by the murine R21D10 Fab (PDB 9AXL; EMD-43969 semiextended, EMD-43983 bent) by Wang et al. [46]. Wang was published after our cohort design and provides an alternative anchor for future docking against the activation-intermediate state. Systematic testing against bent, semiextended, and extended αIIbβ3 ectodomain is deferred to future work.

A related precedent for aptamer-side state-selectivity is Wu et al. [47]: a sub-nanomolar DNA aptamer against CD62L (L-selectin), developed by modified cell-SELEX, that traces and releases CD62L^+^CD8^+^ T cells for CAR T manufacturing. Wu et al. do **not** address integrin αIIbβ3 / β3, but their work makes two relevant points. First, high-affinity SELEX against a cell-state-dependent surface receptor can yield empirically confirmed state-selectivity (a question our in silico cohort cannot address because we model only the β3 monomer in a single conformation). Second, the cell-SELEX templating step (as Teng et al. [28] also do for ITGB3) is an experimental anchor that no current cofolder or restraint-driven docker can substitute for. We therefore treat αIIbβ3-state-selectivity as outside the scope of the present in silico cohort.

**Eighth -- PDB memorisation confound**. PDB memorisation is a known confound for AI cofolders [23]. The cohort includes structures predating the Boltz-2 training cutoff (e.g., 5DO4-H/L, 4DII, 1HUT for thrombin-TBA); Boltz-2 predictions on these are not strictly independent. P01031 / C5 itself has multiple deposited apo and ligand-bound structures predating training cutoffs (e.g., 3CU7 apo C5, 5HCC, 5I5K eculizumab-C5). Because no public Avacincaptad-C5 co-structure was identified as of 2026-05, memorisation of this exact complex from a public PDB deposition is unlikely.

### 4.5 What this paper is *not*

This section pre-empts scope misinterpretation. We explicitly state what this Perspective does *not* claim:

1. **We do not claim that HADDOCK3 is broken for aptamer-protein docking**. The 12-mode catalogue identifies operational failure modes within a specific protocol (blind scale-adaptive AIR on AlphaFold receptors); it does not impugn the underlying physics-based docker. §3.7 demonstrates that Mode 1 (one of the most impactful modes) admits a single-pass mitigation that raises cross-method Jaccard ∼10x without redesigning HADDOCK3.
2. **We do not claim that the blind AIR strategy is unusable**. §3.7 demonstrates the empirical mitigation (pLDDT ≥ 70 prefilter + scale-adaptive N_active) that resolves Mode 1 within the blind-AIR paradigm. The strategy is usable provided the prefilter is used and the limitations in §4.4 and §3.7.7 are taken into account.
3. **We do not claim that the candidate aptamers are non-binders**. The 7/7 positive HADDOCK3 score on P01031 is best characterised as **expected behaviour of the blind top-N % AIR strategy outside the approximate published benchmark coverage region on a 1676 aa multi-domain receptor**. It is not method failure on the aptamers themselves. HADDOCK3 top-1 score sign is not a calibrated binder / non-binder classifier without per-system calibration; it is a function of restraint specification. The 7/7 positive P01031 pattern should not be read as a calibrated no-bind classification (see §3.3 and Figure 2 caption).
4. **We do not claim a statistical threshold for the receptor-size boundary**. The K_D-calibration subset (n = 4 cells) is a deliberately small, matched-condition subset for the Spearman ρ analysis. It should not be conflated with the broader cohort of ≥ 6 biological cognate / intended-cognate cells. The n = 1 P01031 case study is mechanistic, not a generalisation claim.
5. **We do not claim a novel algorithm at §3.7**. The pLDDT prefilter is intuitive in retrospect, consistent with AlphaFold-DB best-practice guidance (pLDDT ≥ 70). The Williams et al. [19] taxonomy (“barbed wire” low-pLDDT class) provides the structural-biology vocabulary, and Holcomb et al. [21] document the protein-ligand analogue of the bias category we extend to the AIR-feed step.
6. **We do not claim a calibrated K_D predictor**. The §3.7.5 n = 4 Spearman ρ shift (-0.80 -> +0.80, exact two-sided p = 0.33) is exploratory cross-method convergence, not significance evidence; it should not be interpreted as predictive performance on prospective targets.
7. **We do not claim universal applicability of the §3.7 mitigation**. §3.7.6 documents a PDB memorisation confound for the Boltz-2 contact reference. The cohort ranges 241-1676 aa with mixed domain architectures. Three classes are outside the present empirical scope or approximate published benchmark coverage region: receptors smaller than ∼100 aa; very-large multi-domain receptors above ∼1500 aa (where Mode 2 / §3.3 applies even with the prefilter); and chemically modified aptamers (PEGylated, XNA, locked nucleic acids).

## 5. Conclusion

We deliver three actionable contributions: a 12-mode failure catalogue collapsed into five broad conceptual classes, a P01031 (1676 aa) case study at the receptor-size boundary, and an empirical Mode 1 mitigation. The mitigation -- a pLDDT ≥ 70 prefilter combined with scale-adaptive N_active = clamp(round(0.08 x N_pLDDT≥70), 15, 80) -- raises cohort blind ⍰ Boltz-2 contact Jaccard ∼10x and corrects a baseline failure that operates ∼40x below random expectation. This work provides an early applicability-domain map for HADDOCK3 + AlphaFold-derived blind AIR on protein-aptamer docking. We adapt the QSAR applicability-domain concept (§4.1) to in silico aptamer screening and encourage similar maps as a baseline reporting practice for new pipelines. Statistical generalisation across the ≥ 1500 aa receptor-size axis remains future work.

## Abbreviations

AIR: ambiguous interaction restraint; a restraint used by HADDOCK to guide docking toward specified interaction regions.
pLDDT: predicted local distance difference test; a per-residue AlphaFold confidence score ranging from 0 to 100. In this study, pLDDT ≥ 70 is treated as high confidence, and pLDDT < 50 as low reliability.
ipTM: interface predicted TM-score; a confidence metric used by AlphaFold-Multimer and Boltz-2 to assess predicted inter-chain interfaces.
iLDDT: interface local distance difference test; a metric for assessing the local accuracy of a predicted interface structure.
cpi_AB: chain-pair interface ipTM between chains A and B; in this study, this refers to the interface ipTM between protein chain A and aptamer chain B.
DockQ: docking pose-quality score; a metric ranging from 0 to 1 for evaluating docking-model quality. Values above 0.23 are commonly treated as acceptable-quality models.
K_D: equilibrium dissociation constant; a lower value indicates stronger binding.
G4 / G-quadruplex: a nucleic acid secondary-structure motif formed by stacked G-tetrads stabilized by monovalent cations such as K^+^.
AF / AF2 / AF3: AlphaFold / AlphaFold 2 / AlphaFold 3 (Google DeepMind structure-prediction models).
RF2NA: RoseTTAFold2NA; a cofolding model for protein-nucleic-acid complexes.
MD: molecular dynamics simulation.
OECD applicability domain: the range of input conditions under which a model or method is considered reliable. In this study, the concept is adapted to define the applicability limits of in silico aptamer screening.
SELEX: Systematic Evolution of Ligands by EXponential enrichment; an in vitro aptamer selection method for identifying nucleic-acid ligands.
CG: coarse-grained representation; a simplified molecular representation that reduces atomistic detail.
scale-adaptive N_active: the number of HADDOCK active residues that scales with receptor size, computed as clamp(round(0.08 x N_pLDDT≥70), 15, 80).

## Data Availability

Cohort data, including 35-pair HADDOCK3 and Boltz-2 results, AIR-generation artefacts, per-pair run directories, aggregate CSV/parquet reports, and figure source data, will be deposited on Zenodo. The DOI will be added when assigned.

## Code Availability

Source code will be released at https://github.com/domy1980/plt_EV_aptamer_agent under the MIT license. The production AIR-generation script is scripts/generate_blind_air_plddt.py; the deprecated historical baseline is scripts/generate_blind_air.py. The HADDOCK3 commit SHA and per-stage provenance metadata will be released with the code and Zenodo archive. Random seeds were not fixed at the driver level, as noted in §2.2.

## Acknowledgements

The author acknowledges AI-assisted editorial feedback during manuscript preparation; all scientific claims, analyses, citations, and conclusions were reviewed and are the responsibility of the author. Computation was performed on Apple M4 Max and Ubuntu 24.04 + RTX A5000 x2 systems. Open-source software is gratefully acknowledged: HADDOCK3 (Bonvin lab, Utrecht), Boltz-2 (Passaro et al., MIT licence), AlphaFold v4 (EMBL-EBI deposit).

## References

1. Giulini M, Reys V, Teixeira JMC, Jiménez-García B, Honorato RV, Kravchenko A, Xu X, Versini R, Engel A, Verhoeven S, Bonvin AMJJ. HADDOCK3: a modular and versatile platform for integrative modelling of biomolecular complexes. J Chem Inf Model 2025;65(13):7315–7324. DOI: 10.1021/acs.jcim.5c00969.

2. Honorato RV, Trellet ME, Jiménez-García B, Schaarschmidt JJ, Giulini M, Reys V, Koukos PI, Rodrigues JPGLM, Karaca E, van Zundert GCP, Roel-Touris J, van Noort CW, Jandová Z, Melquiond ASJ, Bonvin AMJJ. The HADDOCK2.4 web server for integrative modelling of biomolecular complexes. Nature Protocols 2024;19:3219–3241. doi:10.1038/s41596-024-01011-0. (Cited at §1 and §2.2 for the canonical Cβ-neighbor blind-AIR implementation in the HADDOCK web-server tradition.)

3. Versini R, Reys V, Kravchenko A, Honorato RV, Bonvin AMJJ. Integrating the MARTINI2 Coarse-Grained Force Field into HADDOCK3 for Faster Modelling of Large Biomolecular Complexes. bioRxiv 2026;2026.04.25.720800. doi:10.64898/2026.04.25.720800.

4. Xu X, Giulini M, Bonvin AMJJ. Improved prediction of antibody and their complexes with clustered generative modelling ensembles (AlphaFlow + HADDOCK). Bioinformatics Advances 2025;5:vbaf161.

5. Helsem SA, Alfsnes K, Frye SA, Hesselberg Løvestad A, Ambur OH. Two different and robustly modeled DNA binding modes of Competence Protein ComP -- systematic modeling with AlphaFold 3, RoseTTAFold2NA, Chai-1 and re-docking in HADDOCK. PLoS ONE 2025;20(5):e0315160. DOI: 10.1371/journal.pone.0315160.

6. Reys V, Giulini M, Cojocaru V, Engel A, Xu X, Roel-Touris J, et al. Integrative Modeling in the Age of Machine Learning: A Summary of HADDOCK Strategies in CAPRI Rounds 47-55. Proteins 2024 (online 2024-12-30). DOI: 10.1002/prot.26789.

7. Giulini M, Schneider C, Cutting D, Desai N, Deane CM, Bonvin AMJJ. Towards the accurate modelling of antibody-antigen complexes from sequence using machine learning and information-driven docking. Bioinformatics 2024;40(10):btae583. DOI: 10.1093/bioinformatics/btae583.

8. Lam HYI, Pujalte Ojeda S, Brezinova M, Hanke J, Ong XE, Mu Y, Vendruscolo M. Metadiffusion: inference-time meta-energy biasing of biomolecular diffusion models. bioRxiv 2026;2026.02.10.704873. doi:10.64898/2026.02.10.704873.

9. Patel S, Fraser K, Gandavadi D, et al. AptaBLE: A Deep Learning Platform for Aptamer Generation and Analysis. bioRxiv 2026;2026.01.06.698056.

10. Abramson J, Adler J, Dunger J, et al. Accurate structure prediction of biomolecular interactions with AlphaFold 3. Nature 2024;630:493–500. DOI: 10.1038/s41586-024-07487-w.

11. Baek M, McHugh R, Anishchenko I, Jiang H, Baker D, DiMaio F. Accurate prediction of protein-nucleic acid complexes using RoseTTAFoldNA. Nature Methods 2024;21:117–121. DOI: 10.1038/s41592-023-02086-5.

12. Boitreaud J, Dent J, et al. (Chai Discovery Team). Chai-1: Decoding the molecular interactions of life. bioRxiv 2024. DOI: 10.1101/2024.10.10.615955.

13. Bernard C, Postic G, Ghannay S, Tahi F. Has AlphaFold3 achieved success for RNA? Acta Cryst 2025;D81:49–62. DOI: 10.1107/S2059798325000592. (Cited at §1: AF3-RNA benchmark documenting failure to generalise to orphan RNA families without context, INF-nWC < 0.5 across five RNA test sets.)

14. Peng C, Ni W, Liu Q, Hu G, Zheng W. A comprehensive benchmarking of the AlphaFold3 for predicting biomacromolecules and their interactions. Brief Bioinform 2025;26(6):bbaf616.

15. Zhao J, Tram K, Yan H, Li Y. Comprehensive evaluation of artificial intelligence-empowered approaches for protein-aptamer complex prediction. Briefings in Bioinformatics 2026;27(3):bbag206. DOI: 10.1093/bib/bbag206.

16. OECD. Guidance Document on the Validation of (Quantitative) Structure-Activity Relationship [(Q)SAR] Models. OECD Series on Testing and Assessment No. 69, document ENV/JM/MONO(2007)2. OECD Publishing, Paris, 2007. DOI: 10.1787/9789264085442-en. Direct PDF: https://www.oecd.org/content/dam/oecd/en/publications/reports/2014/09/guidance-document-on-the-validation-of-quantitative-structure-activity-relationship-q-sar-models_g1ghcc68/9789264085442-en.pdf (accessed 2026–05-16; verified HTTP 200).

17. Roy K, Kar S, Ambure P. On a simple approach for determining applicability domain of QSAR models. Chemometrics Intell Lab Syst 2015;145:22–29. doi:10.1016/j.chemolab.2015.04.013.

18. Dutschmann TM, Kinzel L, ter Laak A, Baumann K. Large-scale evaluation of k-fold cross-validation ensembles for uncertainty estimation. J Cheminform 2023;15:49. doi:10.1186/s13321-023-00709-9.

19. Williams CJ, Chen VB, Richardson DC, Richardson JS. Categorizing prediction modes within low-pLDDT regions of AlphaFold2 structures: near-predictive, pseudostructure and barbed wire. Acta Crystallogr D Struct Biol 2025;81(10):558–572. doi:10.1107/S2059798325007843. (Cited at §1.1 and §3.7.4 as the structural taxonomy for the failure mechanism.)

20. Yu J, Zhao B, Kurgan L. Comprehensive assessment of AlphaFold’s predictions of secondary structure and solvent accessibility at the amino acid-level in eukaryotic, bacterial and archaeal proteins. Comput Struct Biotechnol J 2025;27:2443–2449. doi:10.1016/j.csbj.2025.05.047. (Cited at §1.1: AlphaFold solvent-accessibility benchmark Pearson r ≈ 0.815 across taxa, supporting the §3.7 pLDDT ≥ 70 prefilter rationale.)

21. Holcomb M, Chang YT, Goodsell DS, Forli S. Evaluation of AlphaFold2 structures as docking targets. Protein Sci 2023;32:e4530. doi:10.1002/pro.4530.

22. Ochoa S, Milam VT. Direct modeling of DNA and RNA aptamers with AlphaFold 3: a promising tool for predicting aptamer structures and aptamer-protein complexes. ACS Synth Biol 2025;14:3049–3064.

23. Škrinjar P, Eberhardt J, Durairaj J, Schwede T. Have protein-ligand co-folding methods moved beyond memorisation? bioRxiv 2025;2025.02.03.636309.

24. Adair BD, Xiong J-P, Yeager M, Arnaout MA. Cryo-EM structures of full-length integrin αIIbβ3 in native lipids. Nat Commun 2023;14:4168. doi:10.1038/s41467-023-39763-0.

25. Jumper J, Evans R, Pritzel A, et al. Highly accurate protein structure prediction with AlphaFold. Nature 2021;596:583–589. (Cited at §2.1 as the AlphaFold 2 architecture used to generate the AlphaFold-DB v4 receptor models.)

26. Russo Krauss I, Spiridonova V, Pica A, Napolitano V, Sica F. Different duplex/quadruplex junctions determine the properties of anti-thrombin aptamers with mixed folding. Nucleic Acids Res 2016;44(2):983–991. doi:10.1093/nar/gkv1384. (Cited at §2.1: RE31 aptamer X-ray crystal with thrombin, PDB 5CMX at 2.99 Å.)

27. Ruckman J, Green LS, Beeson J, et al. 2’-Fluoropyrimidine RNA-based aptamers to the 165-amino acid form of vascular endothelial growth factor (VEGF165). J Biol Chem 1998;273:20556–20567. (Cited at §2.1: Pegaptanib origin; anti-VEGFA SELEX, K_d ≈ 49-130 pM, photo-crosslinking to Cys137.)

28. Teng X, Wang Y, You L, Wei L, Zhang C, Du Y. Screening a DNA aptamer specifically targeting integrin β3 and partially inhibiting tumor cell migration. Anal Chem 2023;95(33):12406–12418. DOI: 10.1021/acs.analchem.3c01995.

29. U.S. Food and Drug Administration. IZERVAY (avacincaptad pegol intravitreal solution) Highlights of Prescribing Information, NDA 217225. Center for Drug Evaluation and Research, U.S. FDA, initial approval 2023-08-04. URL: https://www.accessdata.fda.gov/drugsatfda_docs/label/2023/217225s000lbl.pdf (accessed 2026-05-16). (Reports K_D = 0.69 ± 0.148 nM against human C5; branched 40 kDa PEG modification.)

30. Biesecker G, Dihel L, Enney K, Bendele RA. Derivation of RNA aptamer inhibitors of human complement C5. Immunopharmacology 1999;42:219–230. (Avacincaptad pegol original derivation; ARC1905 / Zimura.)

31. Kuhn LA, Siani MA, Pique ME, Fisher CL, Getzoff ED, Tainer JA. The interdependence of protein surface topography and bound water molecules revealed by surface accessibility and fractal density measures. J Mol Biol 1992;228:13–22.

32. Passaro S, Wohlwend J, et al. Boltz-2: Towards Accurate and Efficient Binding Affinity Prediction. bioRxiv 2025. DOI: 10.1101/2025.06.14.659707.

33. Stafflinger H, Neißner K, Bartsch S, et al. Crystal structure of the class V GTP-binding RNA aptamer bound to its ligand: GTP recognition by a topologically complex intermolecular G-quadruplex. Nucleic Acids Res 2025;53:gkaf1315 (NAR Breakthrough Article). PDB 9HRF, 9HRD, 9HRG.

34. Sun K, Head-Gordon T. KinConfBench: A Curated Benchmark for Cofolding Models on Kinase Conformational States. bioRxiv 2026;2026.04.07.716788.

35. De la O Becerra KI, Brondijk THC, Serna Martin I, Gros P. Structural insights into C3 convertase activity of the classical pathway of complement. Nat Commun 2026;17:993. DOI: 10.1038/s41467-025-67730-4.

36. Jia C, Yang X, Zhao M-H, Tan Y, Xiao J. Complement C3 recognition by C3 convertases. Sci Adv 2026;12:eadz5404. DOI: 10.1126/sciadv.adz5404.

37. Astellas Pharma Inc. Astellas to Highlight New Findings for Geographic Atrophy at ARVO 2026 Annual Meeting. Astellas Newsroom press release, 2026-04-30. URL: https://newsroom.astellas.com/2026-04-30-Astellas-to-Highlight-New-Findings-for-Geographic-Atrophy-at-ARVO-2026-Annual-Meeting (accessed 2026-05-16).

38. Lonfat N, Moreno-Leon L, Lombardi N, Desai A, Khanna H. Nonclinical pharmacology and pharmacokinetic properties of avacincaptad pegol, an aptamer against C5 approved for the treatment of geographic atrophy secondary to age-related macular degeneration. Investigative Ophthalmology & Visual Science (ARVO 2025 Annual Meeting Abstract) 2025;66(8):2539. URL: iovs.arvojournals.org/article.aspx?articleid=2806251.

39. Fukuzawa T, Sampei Z, Haraya K, et al. Long-lasting neutralization of C5 by SKY59, a novel recycling antibody, is a potential therapy for complement-mediated diseases. Sci Rep 2017;7:1080. (Cited at §3.3: PDB 5B71 SKY59-C5 MG1 domain crystal at 2.11 Å, providing a domain-level structural anchor on C5.)

40. King J, Cornwall L, Nica AC, et al. On fine-tuning Boltz-2 for protein-protein affinity prediction. arXiv 2025;2512.06592.

41. Cucchiarini A, Dobrovolná M, Brázda V, Mergny J-L. Analysis of quadruplex propensity of aptamer sequences. Nucleic Acids Res 2025;53:gkaf424. DOI: 10.1093/nar/gkaf424.

42. Bedrat A, Lacroix L, Mergny JL. Re-evaluation of G-quadruplex propensity with G4Hunter. Nucleic Acids Res 2016;44(4):1746–59.

43. Vangone A, Bonvin AMJJ. Contacts-based prediction of binding affinity in protein-protein complexes. eLife 2015;4:e07454. (PRODIGY contact-based predictor.)

44. Krivak R, Hoksza D. P2Rank: machine learning-based tool for rapid and accurate prediction of ligand binding sites from protein structure. J Cheminform 2018;10:39. (Cited at §3.7.1: P2Rank pocket prediction as one of the three method channels in the Jaccard recovery analysis.)

45. Kendall MG, Gibbons JD. Rank Correlation Methods, 5th ed. Edward Arnold, London, 1990. ISBN 0852642628. (Cited at §3.7.5: exact-distribution enumeration of n = 4 Spearman-ρ permutations for the K_D-calibration subset.)

46. Wang L, Wang J, Li J, Walz T, Coller BS. An αIIbβ3 monoclonal antibody traps a semiextended conformation and allosterically inhibits large ligand binding. Blood Adv 2024;8(16):4398–4409. DOI: 10.1182/bloodadvances.2024013177. (R21D10 mAb; PDB 9AXL atomic coordinates of αIIbβ3 ectodomain semiextended conformation; cryo-EM density maps EMD-43969 (semiextended) and EMD-43983 (bent).)

47. Wu AY, Cheng EL, Kacherovsky N, Marking A, Lin-Goldstein A, Heinze CM, Salipante SJ, Jensen MC, Pun SH. Efficient and traceless aptamer-based selection of naïve and early memory CD8 T cells for CAR T cell therapy. Adv Healthc Mater 2026;15(1):e02930. DOI: 10.1002/adhm.202502930. (Sub-nanomolar CD62L DNA aptamer + reversal-agent pair; cell-SELEX-derived; cited here as a recent precedent for aptamer-side selectivity for cell-state-dependent surface receptors, not as an αIIbβ3 / β3 study.)

